# Sample multiplexing for retinal single-cell RNA-sequencing

**DOI:** 10.1101/2024.04.23.589797

**Authors:** Justin Ma, Ting Kuan Chu, Maria Polo Prieto, Yong Park, Yumei Li, Rui Chen, Graeme Mardon, Benjamin J Frankfort, Nicholas M Tran

## Abstract

Rare cell populations can be challenging to characterize using microfluidic single-cell RNA sequencing (scRNA-seq) platforms. Typically, the population of interest must be enriched and pooled from multiple biological specimens for efficient collection. However, these practices preclude the resolution of sample origin together with phenotypic data and are problematic in experiments in which biological or technical variation is expected to be high (e.g., disease models, genetic perturbation screens, or human samples). One solution is sample multiplexing whereby each sample is tagged with a unique sequence barcode that is resolved bioinformatically. We have established a scRNA-seq sample multiplexing pipeline for mouse retinal ganglion cells using cholesterol-modified-oligos and utilized the enhanced precision to investigate cell type distribution and transcriptomic variance across retinal samples. As single cell transcriptomics are becoming more widely used to research development and disease, sample multiplexing represents a useful method to enhance the precision of scRNA-seq analysis.

## INTRODUCTION

Single-cell RNA sequencing (scRNA-seq) is a powerful tool for studying cell type- or state-specific transcription in healthy and diseased tissues^1–3^. Widely used microfluidic droplet-based scRNA-seq platforms enable profiling of transcriptomes from thousands of cells per experiment^4,5^. However, when working with rare or low abundance cell populations, it may not be feasible to obtain the required input cell number for these platforms to be run efficiently. Cells can be pooled from multiple samples for each experiment, but this precludes analysis of gene expression differences across individual samples. This limitation is particularly challenging for experiments where gene expression is expected to differ across biological replicates (e.g., disease models with variable phenotypes, high throughput genetic perturbation screens, or human patient samples). A solution for tracking sample identity in pooled scRNA-seq collections is sample multiplexing, whereby cells from each sample are tagged with unique next generation sequencing (NGS) barcodes prior to pooling^6–10^. Sample barcodes are sequenced in parallel to the transcriptome and sample identities of each cell are extracted bioinformatically. Here, we demonstrate the utility of using sample multiplexing to analyze gene expression patterns of rare cell types.

To establish our sample multiplexing pipeline, we focused on retinal ganglion cells (RGCs), which are the projection neurons that connect the retina to the rest of the brain. We previously generated a cellular atlas of mouse RGC types by scRNA-seq, identifying 46 types^11–13^. Since RGCs account for <1% of all retinal cells, our previous experiments pooled RGCs from ∼6-8 retinas for each scRNA-seq collection, preventing the assessment of sample-specific gene expression^12,14–16^. In this study, we tagged RGCs from individual retinas using cholesterol-modified oligos (CMOs), which utilize a barcode attached to a lipid scaffold that can conjugate to cell surfaces^7^. CMO-labeling efficiently tagged retinal cells and readily integrated with our RGC purification methods. CMO-labeling did not impact transcriptome quality and enabled evaluation of key biological features such as sex-specific differences and assessment of cell-type frequencies in individual retinas. Our analytical pipeline also highlights important considerations for sample de-multiplexing, determination of labeling efficiency, identification of cell multiplets, among other features. Collectively, our studies demonstrate that CMO-labeling enabled the resolution of sample origin in scRNA-seq data. Our research framework optimizes scRNA-seq approaches for studying RGCs in disease models and other experimental conditions and is adaptable to study rare cell populations in other tissues. As scRNA-seq is increasingly being used to evaluate transcriptomic shifts in disease and development, sample multiplexing represents a useful tool which can enhance the precision and interpretability of analysis.

## RESULTS

### CMO-labeling of mouse retinal cells did not impact scRNA-seq quality

To enable tracking of sample identity of RGCs in scRNA-seq collections, we integrated CMO-labeling into our RGC purification protocols^12,13^ (Figure 1A). For each collection, up to six individual retinas from adult *Vglut2-cre;Ai9* mice were enzymatically dissociated and processed in parallel. We modified our RGC purification protocol to incorporate a negative anti-CD73 immunopanning step to deplete rod photoreceptors^17^. Rod-depletion removed ∼60% of the total cells from each sample and reduced the amount of time needed for subsequent fluorescence-activated cell sorting (FACS) steps (Figure S1A-B). Each sample was incubated with a unique CMO barcoding tag and pooled prior to FACS. RGCs (CD90.2+; TdTomato+) were purified by FACS for scRNA-seq (10x Genomics). Consistent with previous experiments, retinal cells maintained high viability (∼77 ± 8.3%) throughout collections and sequenced cells were ∼89 ± 5.1% RGCs (Figure S1C, Table S1). As an additional control, we performed a non-CMO-labeled, rod-depleted pooled RGC collection (Exp0_Unlabeled), which similarly maintained high viability and RGC purity (Table S1).

**Figure 1.**
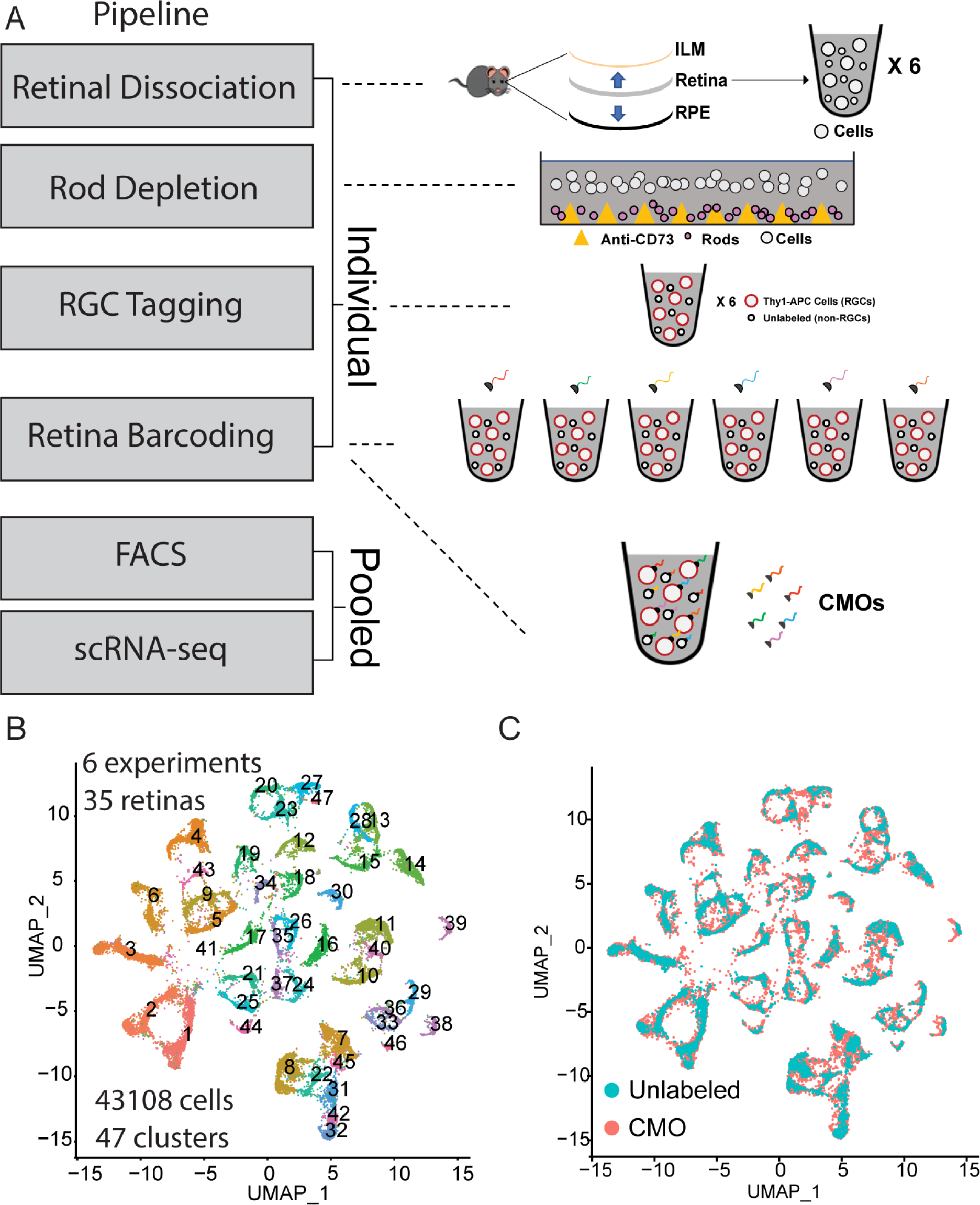
CMO-labeling enables tracking of retinal sample origin in scRNA-seq. (A) Retinas from *Vglut2-Cre;Ai9* mice were enzymatically dissociated and processed in parallel. Rods were depleted by CD73 negative-immunopanning and RGCs were labeled with an anti- CD90.2/Thy1.2 antibody conjugated to APC. Each retina was labeled with a unique CMO barcode and then pooled for RGC enrichment by FACS (TdTomato+;C90.2/Thy1.2+). Collected cells were processed for scRNA-seq. (B) UMAP showing clustering of processed scRNA-seq dataset (6 collections) yielded 43,108 high quality RGC transcriptomes grouped into 47 clusters. This included five CMO-labeled experiments (27 retinas) and one “Unlabeled” experiment (8,734 cells), which was not labeled with CMOs. (C) All clusters contained both CMO-labeled and “Unlabeled” cells, demonstrating transcriptional similarity. ILM: inner limiting membrane; RPE: retinal pigment epithelium. (See also Figure S1 and Table S1).

Overall, we generated 43,108 high-quality RGC transcriptomes by scRNA-seq following quality control (QC) filtering, removal of non-RGCs, and removal of multiplets (Figure 1B). This included 38,801 RGCs collected from 27 CMO-labeled retinas (Table S2). Excluding Exp0 and 1, which had a lower cell input for scRNA-seq, we obtained a mean recovery of 1466 RGCs per retina, ranging from ∼1039 to ∼2219 per retina. Unsupervised clustering identified 47 transcriptionally-related groups. Clustered data recapitulated RGC cell-type specific gene expression patterns previously defined in the reference RGC atlas (Figure S2)^12^. CMO-labeling did not impact clustering as non-CMO-labeled cells co-clustered with CMO-labeled cells (Figure 1C). We detected few significant global gene expression differences between CMO-labeled and Unlabeled cells (Figure S1E). We concluded that incorporation of anti-CD73 immunopanning and CMO-labeling steps did not interfere with RGC purification or impact scRNA-seq data quality.

### Tracking retinal sample origin with CMO-labeling

The sample identity of CMO-labeling retinal cells was determined using computational methods. For each collection, cells were clustered based on detection of their unique CMO barcodes (Figure 2A, Figure S3). We tested two methods to assign sample identity to cells: Cell Ranger Multiplexing Pipeline and the hashtag oligos Demultiplexing package (HTODemux, Seurat)^6,18,19^, (Figure 2B). Both methods distinguish CMO tags from background noise by fitting to a binomial distribution, with the user setting a confidence threshold. We tested multiple thresholds for each method, classifying cells as “Assigned”, Multiplets”, or “Unassigned”, based on the detection of one, multiple, or no CMO barcodes, respectively (Figures 2A, C, Table S2). Assignments largely overlapped between methods, however, HTODemux identified a greater number of multiplets (Figure 2B). For subsequent analyses, we utilized the more stringent HTOdemux assignments. Assignment rate on average was ∼85%, ranging from 70% to 94% across experiments (Figure 2C, Table S2).

**Figure 2.**
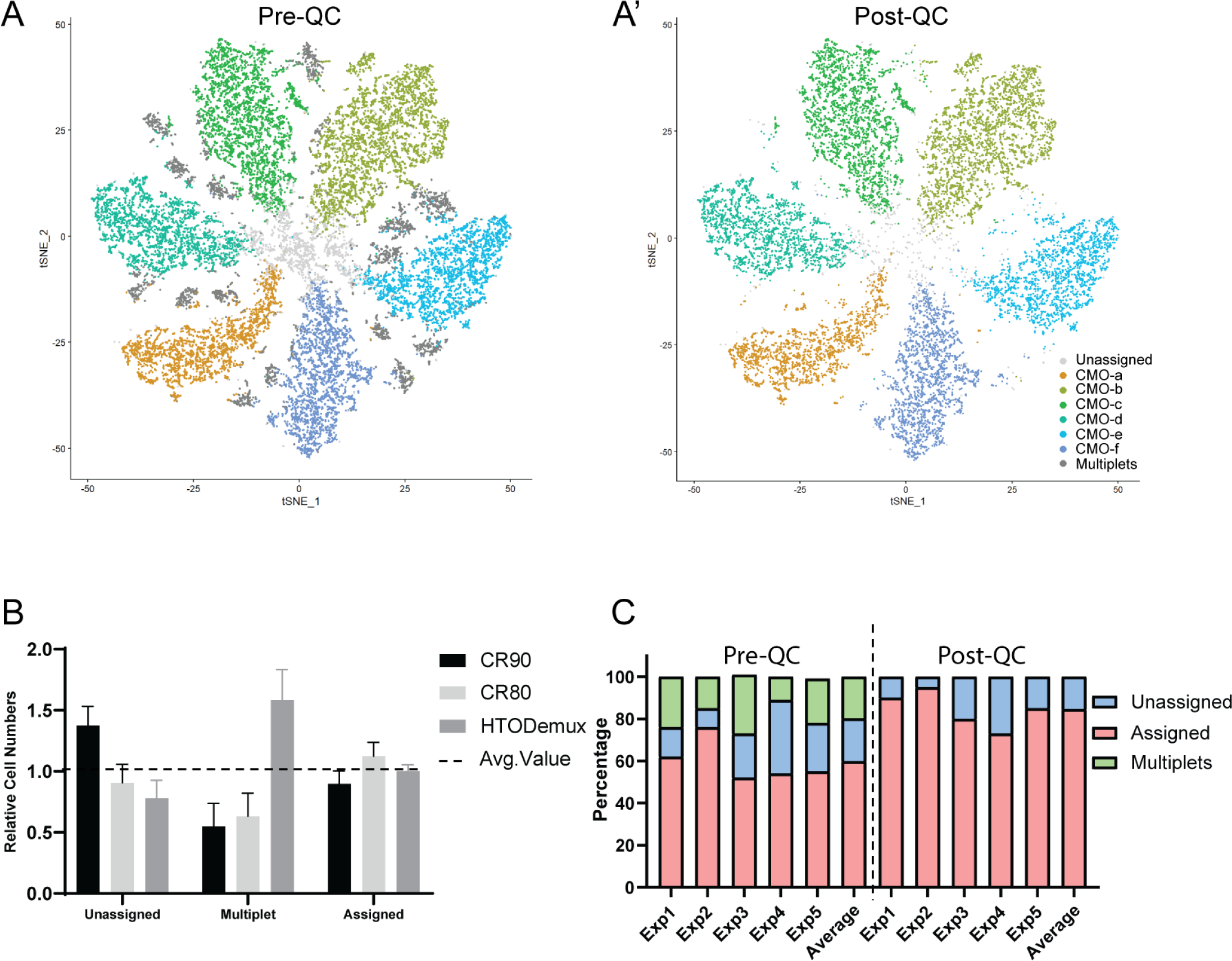
Demultiplexed retinas retain a high percentage of high-quality cells. (A) tSNE showing clustering of cells from Experiment 2 based on CMO-barcode reads. (A’) Multiplets and most ‘Unassigned’ cells were removed by quality control (QC) filters. Resulting clusters largely represent cells labeled with one CMO-barcode, which are predicted to be derived from the same retinal sample. (B) Comparison of the HTODemux assignment method to the Cell Ranger (CR) multiplex pipeline (CR90 and CR80 represent 90% and 80% confidence interval limits used). (C) Comparison of assignment percentages pre- and post- QC filtering. On average, 85 ± 7.7% of the cells that passed QC and multiplet filters were assigned. (See also Figure S3, and Table S2).

As a secondary validation of sample identity assignments, we assessed the expression patterns of sex-related genes in experiments which included CMO-labeled retinas from male and female mice. Expression of genes located on the X (*Xist)* and Y (*Ddx3y*, *Eifs3y*) chromosomes were enriched specifically in cells assigned to samples of the appropriate sex (Figure 3). Multiple other sex-linked genes including *Gm21887* (*Erdr1x*), and *Gm47283* were also significant DEGs (Figure 3A-B). In contrast, the Unlabeled sample, which contained an equal number of male and female retinas, displayed an even distribution of sex-related genes (Figure 3F). It should be noted that the detection of sex-related gene expression varied between experiments (Table S3). While *Xist* was detected in most female-derived cells (95-100%), *Ddx3y* and *Eif2s3y* were detected in fewer male-derived cells (20-70%). A lower level of transcripts of sex-linked genes from the opposite sex were also detected in some cells (Table S3), suggestive of background transcript contamination. Overall, the specific enrichment of appropriate sex-linked genes is indicative that sample assignments were at least qualitatively accurate.

**Figure 3.**
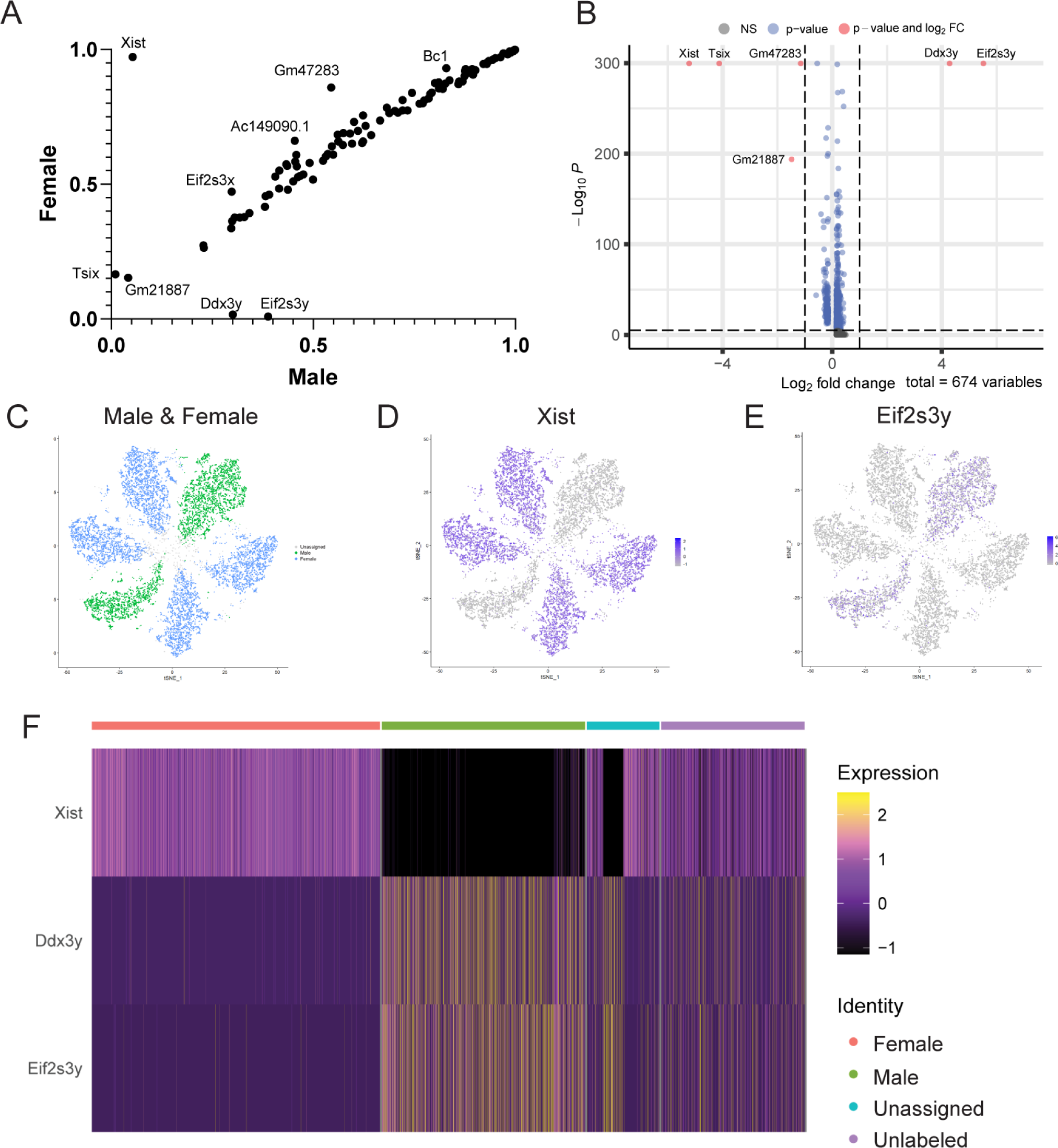
Sample barcoding reveals differential expression of sex-related genes. (A) Top 126 differentially expressed genes (DEGs) comparing cells from male and female samples plotted by percentage of cells with detected transcripts. (B) Volcano plot of top DEGs (FC: fold change, NS: not significant). Six genes had a >1 FC difference between male and female cells, all of which were located on sex chromosomes. (C) tSNE CMO feature plot labeled by sex of originating sample. (D-E) X-linked (*Xist*) and Y-linked (*Eif2s3y*) genes display predicted expression patterns based on sample assignment. (F) Heatmap of all experiments depicting scaled expression of three sex-linked genes segregated by assigned “Male” (green) or “Female” (orange) identity. “Unlabeled Exp0” (purple), which pooled cells from four male and four female mice, was included for comparison. (See also Table S3).

Accurate resolution of multiplets, transcriptional profiles derived from more than one cell, is a common challenge in scRNA-seq analysis. Computational methods to detect multiplets in scRNA-seq data generally rely on features such as transcript count or mRNA heterogeneity, which can be biased if comparing cells with natural differences in these features. We asked if CMO-labeling may improve multiplet detection by comparing HTOdemux assignments to a commonly used computational method, DoubletFinder^20^ (Figure S4). Using a DoubletFinder prediction rate that would yield a similar number of multiplets, we found that the multiplet prediction differed substantially from CMO-defined multiplets. DoubletFinder multiplets were biased towards specific RGC clusters, while CMO-defined multiplets had representation among all clusters (Figure S4B-B’). We concluded that CMO-based detection of multiplets was likely more accurate and removed these multiplets as part of our QC pipeline (Figures 2A’, 2C).

We next assessed RGC type distributions across CMO-labeled retinas. We overlayed sample assignments onto our clustered transcriptomic data (Figure 4A). We found that most clusters contained cells from every CMO-labeled retina at relatively even distributions, with the exceptions being relatively sparse RGC types. (Table S4). “Unassigned” cells, for which sample identity was not assigned, demonstrated a similar distribution pattern (Figure 4A), suggesting these cells were qualitatively similar to “CMO-Assigned” cells. RGC frequencies also tracked closely with distributions in the RGC atlas^12^ and Unlabeled cells (Figure 4B). These results demonstrated that CMO-labeling does not bias collection towards certain RGC types and that RGC types exhibit predictable frequencies across biological replicates.

**Figure 4.**
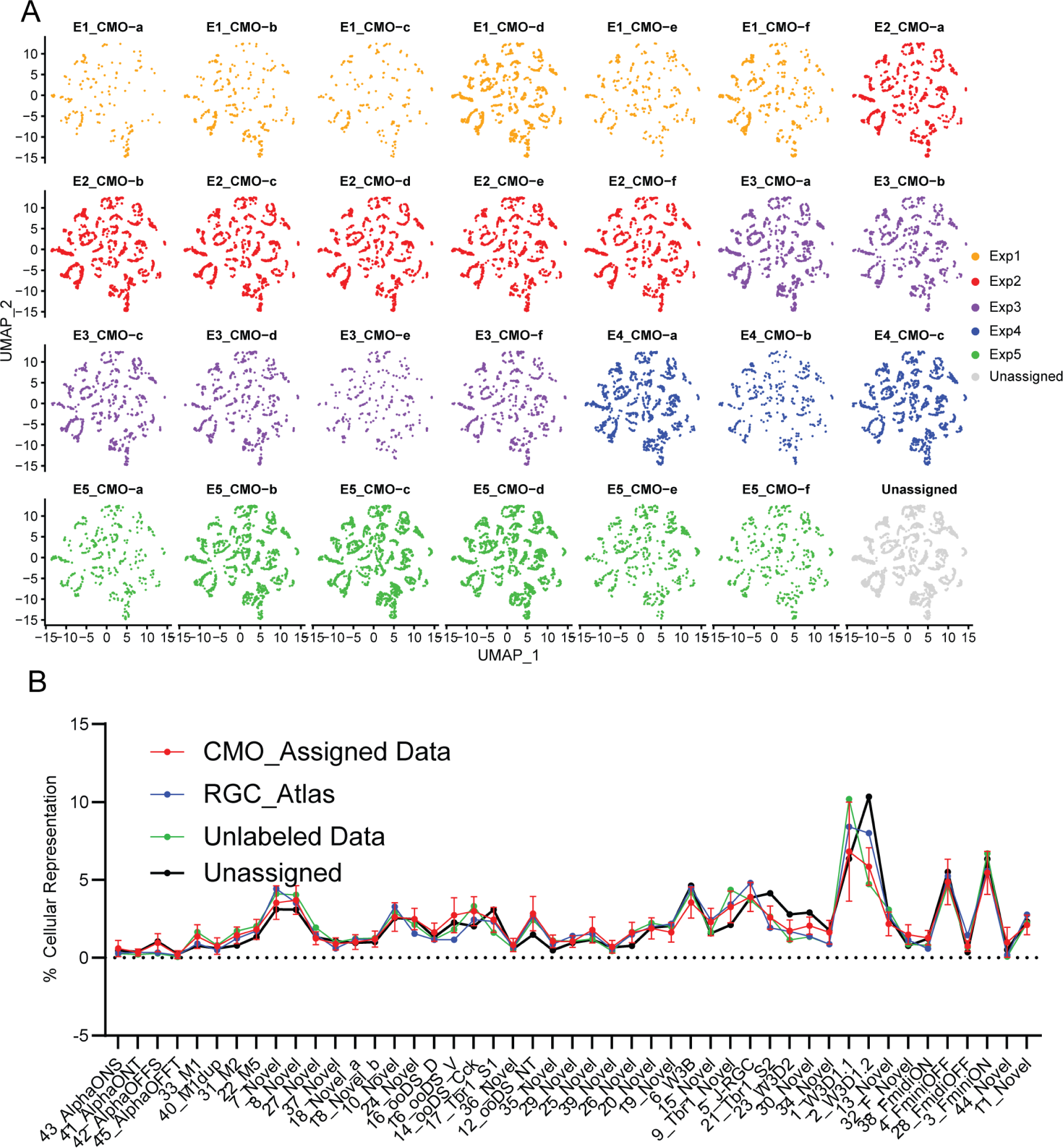
RGC type distribution is consistent across CMO-labeled retinas. (A) UMAPs showing distribution of RGCs from individual retinas across transcriptomic clusters (Exp0 not included). RGCs collected across CMO-labeled samples were evenly distributed across clusters. Unassigned RGCs were also distributed across clusters. (B) Line plot depicting RGC type percent representation between the “RGC_Atlas” (Tran et al. 2019^12^), “Unlabeled Data” (Exp0), “CMO-labeled Data” (N = 27 retinas), and the Unassigned data. This graph indicates RGC type distribution was consistent across samples, labeling method, and with previously published datasets. (See also Table S4).

### RGC gene expression across samples and cells

To determine whether cells from different retinas display gene expression differences, we analyzed global expression patterns on a per-retina basis (Figure 5). Comparison of gene expression from cells assigned to different retinas within each collection were highly similar (> 0.97 Pearson correlation), even though each retina was dissected and processed separately (Figure 5). Cells from retinas collected in different experiments were also highly correlated (>.87 Pearson correlation) but displayed more gene variance, which was likely due to batch effects (Figure 5C). We compared transcriptomic profiles between RGCs derived from left or right eyes from the same animal and detected no significant DEGs (Figure S5). Consistent with our analysis of sex-related gene expression in male and female samples, sex-linked genes were among the most variable genes across samples (Figure 5D-E, Table S5). There were additional genes such as *Lars2* or *Gm42418* with substantial levels of variation but no correlated phenotypic distinction. In general, global expression patterns among retinal samples were consistent within and between experiments. In addition, CMO-labeling enabled the resolution of sex-specific differences, a valuable feature of the experimental design.

**Figure 5.**
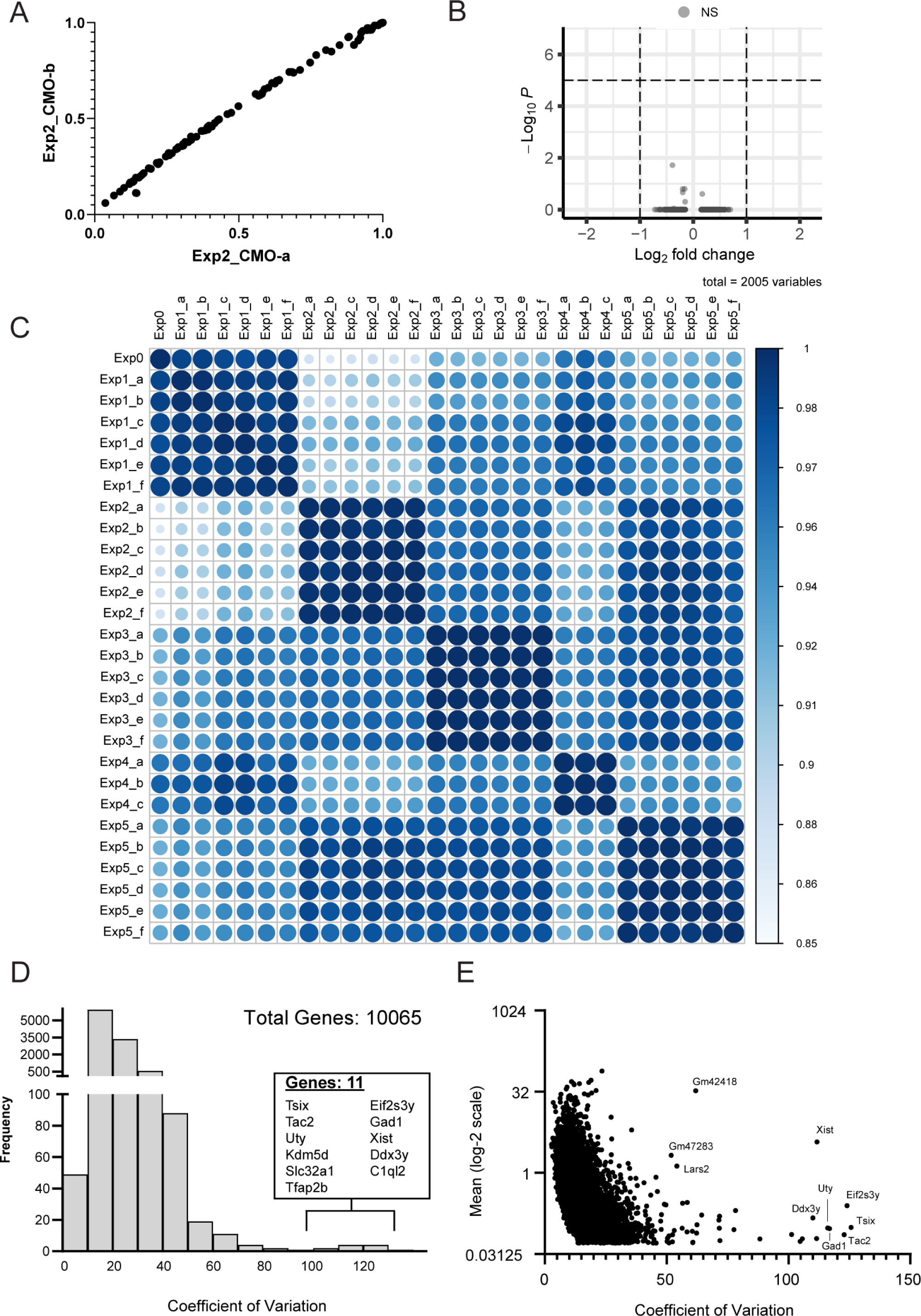
Gene expression patterns are highly correlated between CMO-labeled retinas. (A-B) A representative comparison showing the low transcriptional variance between cells assigned to different CMOs within each collection. No significantly different genes were identified comparing cells assigned to CMO-a and CMO-b from Exp2. (C) Correlation analysis (Pearson) comparing the average gene expression of the top 5000 variable genes between all retinas. Correlations ranged from 0.87-1.00 across all samples and 0.99-1.00 comparing samples from the same experiment, indicating high transcriptional similarity among cells derived from different retinas within and between experiments. D) Histogram showing distribution of the CV across all detected genes (scaled mean >0.05). The 11 most variable genes (>100 CV) are listed. (E) Scatterplot comparing CV to mean expression level. (See also Figure S6 and Table S5, S6).

We next assessed whether type-specific gene expression patterns were consistently expressed across retinal samples. We analyzed the variance in gene expression of the type-defining markers (defined in Table1, Figure S7A) between retinas and assessed whether variance was associated with cluster abundance (Figure S6, Table S6). We found that variance in marker expression and cluster abundance were generally negatively correlated. Higher abundance types such as F_miniON and W3D1.1 had the lowest variability, while scarce types such as AlphaOFFT and AlphaONT were among the highest. Most types, including low abundance populations, recapitulated type-specific expression patterns across retinas with some specific exceptions (Table S6). A notable outlier was 44_Novel, which was not well resolved in our dataset. This indicates that the chosen markers robustly identify RGC types, even on an individual retina basis. Overall, we demonstrate that CMO-labeling enabled assessment of RGC type-specific gene expression in individual retinas but highlights a limitation for interrogation of expression patterns in low abundance types.

### RGC Atlas version 1.1

While not the primary objective of this study, our expanded RGC scRNA-seq dataset enabled a re-examination of RGC type annotations and expression patterns. The 47 clusters identified in our analysis correlated directly with the RGC atlas types with two exceptions. 16_ooDS_DV (on-off direction selective dorsal and ventral types) and 18_Novel each split into two clusters that we annotated as: 16_ooDS_D, 16_ooDS_V, 18_Novel_a, and 18_Novel_b. The division of 16_ooDS_DV is consistent with Tran et al. 2019^12^, which determined this group contained two transcriptionally similar types that could be manually subdivided based on the expression of *Calb1* and *Calb2,* established markers for the ooDS_D and ooDS_V types, respectively^21^. Indeed, the ooDS_D and ooDS_V clusters were specifically enriched for *Calb1* or *Calb2* (Figure 6B-B’’). 18_Novel_a and 18_Novel_b were distinguished by the expression of *Pcdh11x* and *Pcdh20/Prkg2*, among other markers (Figure 6C-C’’, Figure S7, Table 1). 18_Novel, along with other ‘Novel’ types described in Tran et al. 2019^12^, were molecularly defined but have not been completely characterized. However, annotation of these four types was independently verified in a large-scale integrated analysis of mouse retinal scRNA-seq datasets^22^. In addition, we analyzed our clustered dataset to validate and revise cell-type specific markers. We identified markers that designate transcriptomically-related groups of RGCs including groups defined by the expression of specific transcription factors like *Irx3*, *Bnc2*, or co-expression of *Mafb* and *Kcnd2*, respectively (Table 1, Fig S7). We defined a minimal set of molecular markers that identify each type, requiring no more than five markers for each (Table 1). The combined expression of our updated markers was highly type-specific and sufficient to resolve all adult RGC types (Figure S7). Lastly, we curated annotations from two recent studies which used Patch-seq to associate molecular types with functional characteristics (Table 1)^23,24^. Our updated RGC annotations improved type resolution and provide useful information for identifying or gaining genetic access to specific types.

**Figure 6.**
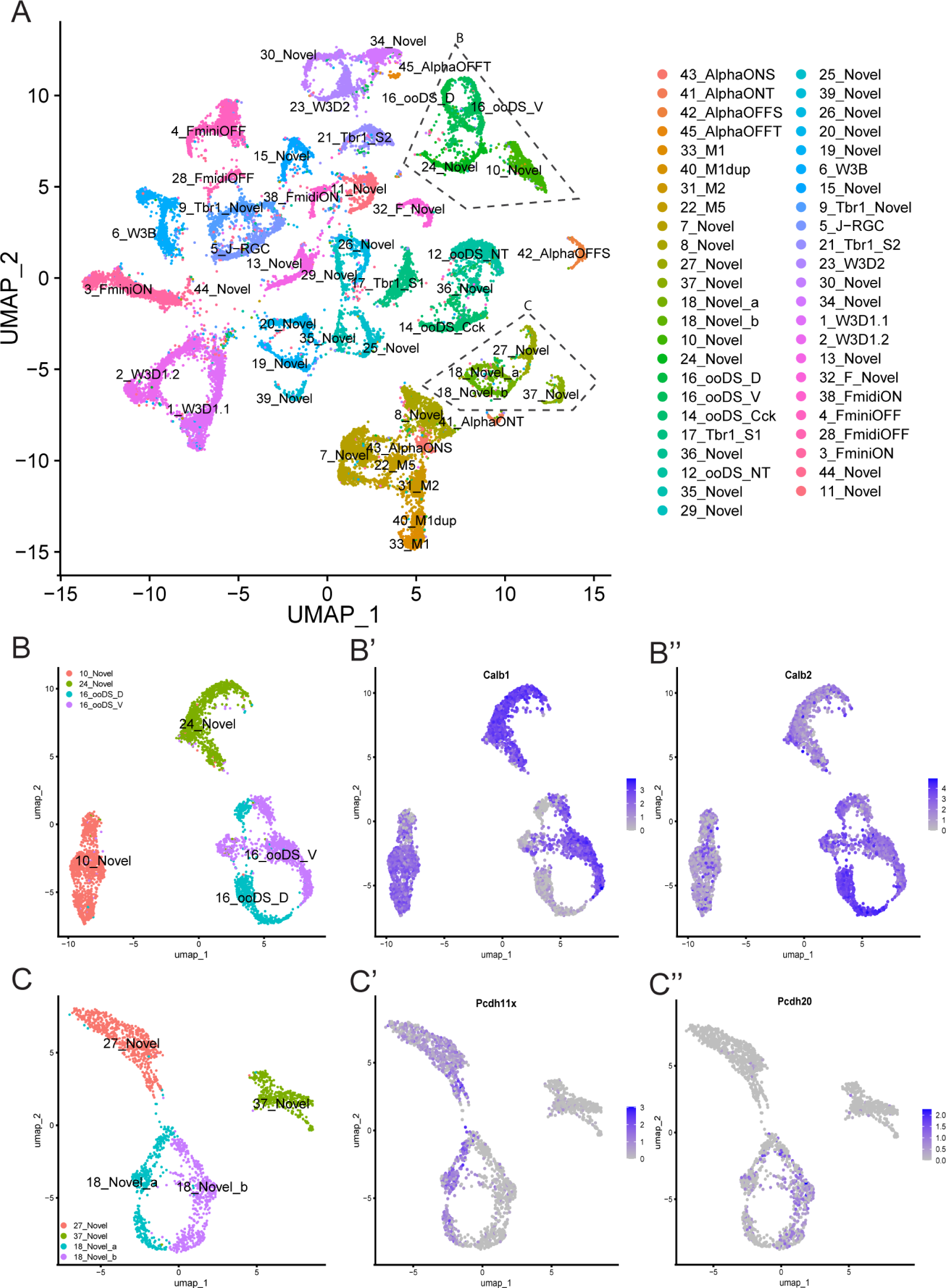
Refinement of the RGC Atlas annotation. (A) UMAP of the transcriptomic data with revised labels. Clusters highlighted by dashed lines are further examined in (B). (B) Featureplots depicting enrichment of *Calb1* and *Calb2* expression in ooDS_V and ooDS_D, respectively. (C) Featureplots depicting enrichment of *Pcdh11x* and *Pcdh20* expression in 18_Novel_a and 18_Novel_b, respectively. (See also Figure S7).

**Table 1.** Retinal Ganglion Cell Reclassification.

| Size Order | Naming |  | Markers | Functional Studies |  |  |
| --- | --- | --- | --- | --- | --- | --- |
| Cluster # | Name | Atlas_name | Type_Specific_Markers | Huang 2022 (blue)<sup>a</sup> | Goetz 2022 Label | Goetz 2022 Function |
| 45 | 43_AlphaONS | 43_AlphaONS | Spp1 Opn4 Il1rapl2 | ON | ON alpha | ON sustained |
| 46 | 41_AlphaONT | 41_AlphaONT | Spp1 Kit |  |  |  |
| 39 | 42_AlphaOFFS | 42_AlphaOFFS | Spp1 Fes | OFF |  |  |
| 47 | 45_AlphaOFFT | 45_AlphaOFFT | Spp1 Chrm1 Tpbg | OFF | OFF tr alpha | OFF transient |
| 32 | 33_M1 | 33_M1 | Opn4 Adra2a | ON | M1 | ON sustained |
| 42 | 40_M1dup | 40_M1dup | Opn4 Nmb |  |  |  |
| 31 | 31_M2 | 31_M2 | Opn4 Pnoc | ON |  |  |
| 22 | 22_M5 | 22_M5 | Opn4 Gpc5 Cdhr1 |  |  |  |
| 8 | 7_Novel | 7_Novel | Eomes Cdh6 Gm11100 |  |  |  |
| 7 | 8_Novel | 8_Novel | Eomes Cdh6 Irx1 | ON |  |  |
| 29 | 27_Novel | 27_Novel | Irx3 Prkcg | ON | ONhOS SmRF | ON OS |
| 38 | 37_Novel | 37_Novel | Irx3 Ceacam10 | ON-OFF |  |  |
| 36 | 18_Novel_a | 18_Novel | Irx3 Pcdh20 Pcdh11x |  | ON bursty | SbC/Other |
| 33 | 18_Novel_b | 18_Novel | Irx3 Pcdh20 Prkg2 |  | ON bursty | SbC/Other |
| 14 | 10_Novel | 10_Novel | Bnc2 Gpr88 Pde11a | ON | ONDS | DS |
| 15 | 24_Novel | 24_Novel | Bnc2 Gpr101 Tafa4 Ptprt | ON-OFF | ooDS T | DS |
| 28 | 16_ooDS_D | 16_ooDS DV | Bnc2 Calb1 Col25a1 Serpinb8 Efnb2 | ON-OFF | OODS D | DS |
| 13 | 16_ooDS_V | 16_ooDS DV | Bnc2 Col25a1 Fam129a | ON-OFF | OODS D | DS |
| 10 | 14_ooDS_Cck | 14_ooDS_Cck | Dab1 Cck Tent5a | ON | ON delayed | SbC/Other |
| 16 | 17_Tbr1_S1 | 17_Tbr1_S1 | Tbr1 Slc35f3 Irx4 | ON-OFF |  |  |
| 40 | 36_Novel | 36_Novel | Dab1 Coch Stxbp6 Cd44 |  | ONhOS LgRF | ON OS |
| 11 | 12_ooDS_NT | 12_ooDS_NT | Neurod2 Satb2 Mmp17 | ON-OFF |  |  |
| 37 | 35_Novel | 35_Novel | Neurod2 Satb2 Chrm2 Cidea |  |  |  |
| 35 | 29_Novel | 29_Novel | Neurod2 Satb2 Cpne5 |  |  |  |
| 24 | 25_Novel | 25_Novel | Neurod2 Satb2 Slc17a7 | ON-OFF | bSbC | SbC/Other |
| 44 | 39_Novel | 39_Novel | Neurod2 Satb2 S100b |  |  |  |
| 26 | 26_Novel | 26_Novel | Neurod2 Satb2 Penk Aga |  | sSbC EW27 | SbC/Other |
| 21 | 20_Novel | 20_Novel | Neurod2 Satb2 Penk Prdm8 Gal | OFF |  |  |
| 25 | 19_Novel | 19_Novel | Neurod2 Satb2 Penk Prdm8 Ipcef1 | OFF |  |  |
| 6 | 6_W3B | 6_W3B | Penk Zic1 | ON-OFF | HD2 | ON OFF small RF |
| 19 | 15_Novel | 15_Novel | Penk Meis2 Col11a1 | ON-OFF |  |  |
| 9 | 9_Tbr1_Novel | 9_Tbr1_Novel | Tbr1 Slc7a11 Satb2 | OFF | OFFhOS | OFF sustained |
| 5 | 5_J-RGC | 5_J-RGC | Tbr1 Pcdh20 | OFF | OFFvOS | OFF sustained |
| 12 | 21_Tbr1_S2 | 21_Tbr1_S2 | Tbr1 Calca Zeb2 | OFF | OFF tr SmRF | OFF transient |
| 23 | 23_W3D2 | 23_W3D2 | Mafb Kcnd2 Prokr1 |  |  |  |
| 20 | 30_Novel | 30_Novel | Mafb Kcnd2 Vgf | ON | ON tr SmRF | ON transient |
| 27 | 34_Novel | 34_Novel | Mafb Kcnd2 Fzd6 Tpbg | ON |  |  |
| 1 | 1_W3D1.1 | 1_W3D1.1 | Trarg1 Amigo2 Cmtm8 | OFF |  |  |
| 2 | 2_W3D1.2 | 2_W3D1.2 | Trarg1 Lypd1 Ldb2 |  | UHD | ON OFF small RF |
| 17 | 13_Novel | 13_Novel | Trarg1 Runx1 Ntrk1 | ON-OFF | HD1 | ON OFF small RF |
| 30 | 32_F_Novel | 32_F_Novel | Foxp2 Coch |  |  |  |
| 34 | 38_FmidiON | 38_FmidiON | Foxp2 Anxa3 |  | ONvOS SmRF | ON OS |
| 4 | 4_FminiOFF | 4_FminiOFF | Foxp2 Gria1 |  |  |  |
| 43 | 28_FmidiOFF | 28_FmidiOFF | Foxp2 Sema5a |  |  |  |
| 3 | 3_FminiON | 3_FminiON | Foxp2 Irx4 | ON-OFF | F-mini-ON | ON OFF small RF |
| 41 | 44_Novel | 44_Novel | Bhlhe22 Fxyd6 | ON |  |  |
| 18 | 11_Novel | 11_Novel | Cacna1e Gm17750 | ON-OFF | LED | ON OFF small RF |

a. Assigned function is based on the responses to blue light

## DISCUSSION

We have developed an RGC scRNA-seq protocol that incorporates sample multiplexing using CMO-based barcoding. Multiplexing did not affect data quality, cell type prediction, or expression patterns. To our knowledge, this is the first scRNA-seq study to evaluate RGC cell type distributions and expression patterns from individual retinas. Sample multiplexing has several benefits for studying gene expression in rare cell populations including increasing the number of biological replicates and directly matching transcriptomic profiles with sample phenotypes and other metadata like sex or age. Implementation of these approaches enhances the efficiency and scalability of testing different experimental conditions. It is important to note that there are several methods for sample multiplexing for scRNA-seq including antibody-based methods such as Cite-seq^6^, or methods which target the genome such as CellTag Indexing^9^. We selected the CMO-labeling approach for this study because this method is readily integrated with our collection protocols. However, the optimal multiplexing approach will differ by experiment.

Using CMO-based multiplexing, we demonstrated a sample assignment rate of ∼85%, ranging from 70-94% across samples, with an average recovery of 1466 RGCs/retina post-quality control. Subjectively, we found that assignment rate may correlate with sample quality, as samples in which more cells were filtered out by standard QC metrics also had the most unassigned cells. Within genotypically homogenous samples like tissues from congenic mice, it is challenging to independently verify classification accuracy. We utilized sex-related genes to estimate cross-sample ‘contamination’ on a qualitative level. In general, appropriate enrichment of sex-specific genes was observed across samples but notably the detection level (e.g., percent of cells expressing each sex-linked marker) and background contamination was variable across collections. While informative, evaluation of sex-related genes only divided samples into two groups for each collection, whereas six retinas were assessed. Therefore, additional sample-specific features would be required to quantitatively evaluate cross-sample contamination. Overall, we conclude CMO-labeling efficiently and accurately identified retinal origin for most cells.

An additional useful feature of CMO-labeling was the ability to accurately identify multiplets. Multiplets are present in most scRNA-seq datasets and can impact clustering and confound interpretation, if not properly resolved^25–27^. Bioinformatic methods are commonly used to identify multiplets within scRNA-seq data^20,28,29^, however, these rely on gross transcriptomic metrics such as transcripts or UMIs per cell, which can differ by cell type. Without a secondary method, it is difficult to verify the accuracy of computationally-derived multiplet detection. Our analysis demonstrated that multiplet detection differed substantially between CMO-based assignments and the widely used computational method DoubletFinder^20^. CMO-based multiplet assignments were more evenly spread across RGC types, suggesting less type-specific bias. A limitation to CMO multiplet assignments is that precision inherently scales with the number of samples labeled. For example, our experiments assessed 6 CMO-labeled retinas per experiment, so theoretically ∼83% of multiplets would be expected to have two or more different CMO tags, while ∼17% would have the same tag. We conclude that sample multiplexing improved detection of multiplets but that it may be advisable to consider additional computation methods for independent verification.

The primary purpose of incorporating sample multiplexing into scRNA-seq pipelines is to resolve type-specific transcriptional differences in individual samples. Global and type-specific expression patterns were highly consistent across samples. We found little expression differences when comparing cells from ‘left’ and ‘right’ eyes. In contrast, sex-related genes demonstrated appropriate enrichment patterns based on predicted sample origin. Type-specific markers were generally consistent across samples, but these patterns were less-well resolved for ‘low abundance’ types. The top 5 scarcest clusters had an average of 5.4 cells/retina, 5 median clusters had an average of 19.8 cells/retina, and the top 5 most abundant clusters had an average of 57.2 cells/retina. Thus, appropriate caution should be used in the assessment of ‘low abundance’ types in individual samples. Our results also demonstrated that RGC types were evenly distributed across samples, supporting the conclusion that these molecular groupings represent cell types, rather than cell states or batch-specific groupings. Overall, type-specific expression differences across retinas were readily detectable. As scRNA-seq is increasingly used to evaluate transcriptomic shifts in disease and development, sample multiplexing is a useful tool to improve the precision of scRNA-seq analysis and enhance testing across experimental conditions and biological replicates, particularly for rare cell populations.

## LIMITATIONS OF THE STUDY

Our study concludes that CMO-based multiplexing enhances precision of scRNA-seq with few drawbacks. However, there are limits to its applicability. The labeling efficiency can be variable and it may be difficult to empirically determine the precise accuracy and efficiency of labeling in an experiment. Identification of multiplets is improved by analysis of CMO barcodes but remains imperfect, though this was partially circumvented by focusing on multiplet clusters rather than individual cells. In addition, despite enabling resolution of RGCs across individual samples, the capacity to assess relatively low abundance RGC types on a per retina basis is limited, as demonstrated by the increased variance in marker gene expression.

## Acknowledgements

This work was performed using the support of these funding sources: R00EY029360, Whitehall Foundation, TIRR Mission Connect to N.M.T., EY025601, EY033458, Research to Prevent Blindness (Unrestricted Grant), Retina Research Foundation. In addition, this work was performed at the Single Cell Genomics Core at Baylor College of Medicine partially supported by NIH S10OD025240 and CPRIT RP200504.

## Author Contributions

All authors have read and approved the final version of this manuscript and meet the Journal’s criteria for authorship. All experiments were performed in the laboratories of N.M.T. B.J.F., G.M. and R.C. by J.M., N.M.T., T.K., M.P., Y.P., and Y.L. The study was conceived and designed by J.M., Y.P., N.M.T., B.J.F. and G.M.. Data was analyzed and interpreted by J.M., N.M.T., B.J.F, and G.M.. The manuscript was drafted and revised by J.M., N.M.T., B.J.F, and G.M..

## Supplemental Information

Document S1. Figures S1-S6 and Tables S1-S6

## Methods

### Resource Availability

#### Lead contact

Any additional information required to reanalyze the data reported in this paper is available from the lead contact upon request. Further information and requests for resources or reagents should be directed to and fulfilled by Nicholas M Tran.

#### Materials Availability

This study did not generate any new reagents.

#### Data and Code Availability

Single-cell RNA-seq data (raw and processed) have been deposited to the gene expression omnibus along with the appropriate metadata and are publicly available at the time of publication. Accession number is listed in the key resources table. All original relevant code is stored as a GitHub repository and is publicly available. Hyperlink is available in the key resources table.

### Experimental Model: Mouse Lines

We used *Vglut2-Cre* (Jax# 028863,^30^) mice mated to the Cre-dependent *Ai9-Tdtomato* (Jax# 007909,^31^) reporter line, to fluorescently label RGCs. All animals were treated in accordance with the National Institutes of Health (NIH) guidelines, the Association for Research in Vision and Ophthalmology (ARVO) Statement for Use of Animals in Ophthalmic and Vision Research, and the Baylor College of Medicine Institutional Animal Care and Use Committee (IACUC) welfare guidelines. All experiments are carried out using mice from 7-12 weeks of age with equal inclusion of both sexes.

### Method Details

#### Cell Preparation and Sequencing

##### Retinal Dissociation, RGC enrichment, and CMO labeling

Retinas were dissociated as previously described with some modifications^12,13,17^. Mice were euthanized and retinas were dissected out in oxygenated Ames media in the presence of Actinomycin D (ActD, 30 µM) along with the removal of ciliary bodies, vitreous, and inner limiting membranes (ILM). Up to six retinas were used per experiment and processed independently. Retinas were digested in papain and dissociated into single cells by gentle manual trituration using a pipette in ovomucoid solution.

Dissociated retinal cells were negative-immunopanned using anti-mouse CD73 (#550738; BD Pharmingen, San Jose, CA; 0.20% diluted in 50mM Tris-HCl pH 9.5) to deplete rod photoreceptors. The remaining cells are resuspended in our working cell buffer solution (AMES/0.4% BSA + 3 µM ActD) and incubated with CD90.2/Thy1.2 antibody conjugated to APC (#17-0902-83; Invitrogen, Carlsbad, CA; 1 μl per 5 million cells) to label RGCs. Cells were then washed and labeled with CMO’s using the 10x Next GEM Single Cell 3’ Reagent Kit v3.1 with Feature Barcoding Technology for Cell Multiplexing (PN-100268, PN-1000261, PN-1000243).

Each retina-derived cell suspension is mixed with a unique CMO at baseline concentration and incubated for exactly five minutes and subsequently the reaction is paused on ice. Each cell suspension was then washed by resuspension in AMES/1% BSA + 3 µM ActD followed by centrifugation. CMO-labeled samples were pooled, counterstained with DAPI to identify non-viable cells, and sorted using FACS. RGCs were purified by FACS based on co-expression of CD90.2/Thy1.2 (APC) and Vglut2 (TdTomato).

##### Single-cell Sequencing

Cells are resuspended in Ames/0.4% BSA and triturated into a single-cell suspension for library construction as per the manufacturer’s protocol for the 10x genomics chromium platform. Library construction is performed separately for gene expression and CMOs. Approximately 10,000-25,000 cells were loaded per channel. Libraries were sequenced using the Illumina Novaseq 6000 with paired-end sequencing (GEX: TruSeq Read1, TruSeq Read2; CMO: Nextera Read1, Nextera Read2).

#### Bioinformatics

##### Cell Ranger

Sequencing Alignment was done using the Cell Ranger v3.1 mkfastq and multi pipelines. Cell Ranger based demultiplexing was performed but was not the primary demultiplexing method used for assessment of cell to retina assignment. Sequences were aligned to the GRGCm38/mm10-2020A genome reference build to generate cell-by-gene and cell-by-CMO matrices for gene expression and CMO multiplexing, respectively.

##### Computational Methods to Define RGC Collection

Gene expression matrices were processed using Seurat 4.0 in R-programming language. Our cumulative RGC dataset includes six collections (Exp0 was Unlabeled, Exp1-5 were CMO-labeled). The expression matrices were integrated using “Harmony”^32^, followed by a series of QC filters resulting in a total RGC number of 43,108 cells. Data was processed as follows: the data was first normalized and scaled using the functions of the Seurat package^18^. Then, randomized PCA analysis was used for dimensional reduction accounting for 100 principal components and batch correction was performed using the Harmony algorithm. Then, we performed pattern classification by calculating the k-nearest neighbors graph (elucidean distance) followed by clustering using the Louvain algorithm (k. param = 30). For quality control (QC), we removed non-RGCs based on marker expression, cells with poor quality transcriptomes (mitochondrial genes > 10%, UMI <= 2400, features <= 1200), and multiplets as defined by CMO classification.

##### Batch Correction

We used “Harmony” to align samples for UMAP clustering. We separately performed additional batch-correction using the R package “batchelor”^33^, specifically the "rescaleBatches” function to correct the expression data between samples for the purposes of differential gene analysis.

##### Computational Methods for Demultiplexing

From Cell Ranger, the CMO multiplex pipeline generated a CMO counts matrix as well as assignment-confidence levels calculating using Cell Ranger’s tag assignment algorithm. The CMO counts matrix was similar to the traditional gene expression matrix, but the features were the CMO counts per cell. We utilized two methods to assign CMO labels, Cell Ranger (10x Genomics) and HTOdemux^6^. Each method was used to identify cells as assigned (labeled with one CMO), blank (no CMO label), or a multiplet (two or more CMO labels). Cell Ranger’s tag assignment algorithm was based on a customized Expectation-Maximization algorithm^19,34^ to fit a binomial distribution. The algorithm assumes a gaussian distribution of counts containing background noise and actual counts from cell tags. Cells with a confidence threshold >0.9 were assigned. The HTODemux algorithm utilizes k-medoid clustering on the normalized CMO counts and fits a negative binomial distribution for each CMO. Barcodes with CMO signals >0.99 quantile was considered “positive” for that CMO. As depicted in Figure 2B, we found that this method was more stringent in classifying multiplets in our dataset. For cluster visualization, the CMO counts matrix was processed as a Seurat object following a standard workflow and visualized by tSNE distribution.

##### Multiplet Comparison

For comparing DoubletFinder and CMO-based multiplets, we analyzed data from Exp2 in isolation. This analysis is performed on an isolated sample to reduce potential variables and this collection was selected because it contained a large number of high-quality cells and a high efficiency of CMO-labeling. Prior to multiplet comparison, all QC filters were applied with the exception of multiplet removal. DoubletFinder was performed using the associated R package^20^ and using sctransform for pre-processing, accounting for 50 principal components, and assumed a 15% doublet rate.

##### DEG Analysis

Top DEGs are determined by using the FindMarkers function as part of the R Seurat package using the “bimod” test^35^. Percentage of cells expressing each gene and fold change within expressing cells were both used in determination of DEGs. Retina-wide comparisons were evaluated using averaged expression levels (for each gene, expression was averaged across all cells per retina). Similarity of pseudo-bulk retinal transcriptomes was assessed using Pearson correlation between the average expression of the top 5000 variable genes between all 27 CMO-labeled retinas. Gene variance was assessed by calculating the coefficient of variation (CV, standard deviation divided by the mean) of each gene between all 27 retinas. For top gene variance rankings, we removed lowly expressed genes (scaled average <0.05) of which CV calculations may be inaccurate. Cluster-specific gene variance is similarly assessed using CV but applied for each cluster on only the associated marker genes. For cluster-specific variance calculations, retinas from Exp1 were not included because of low cell representation and retinas without cells belonging to a particular cluster were not factored in for the data of that cluster.

### Resources Table

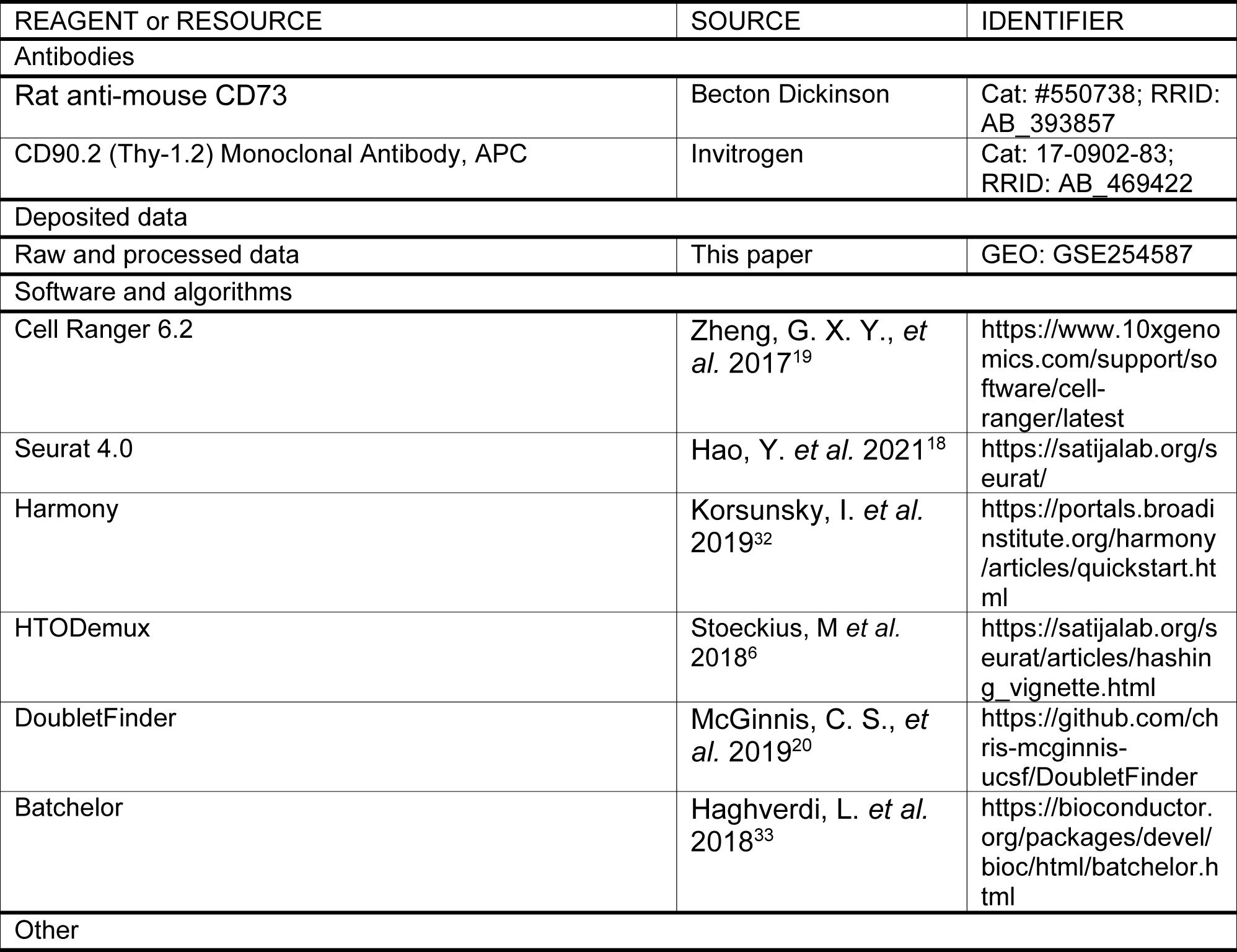
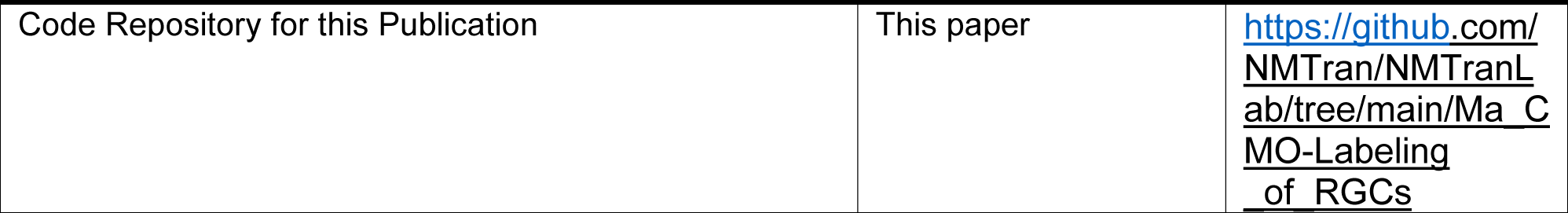

