## Supplemental Figures and Tables for "Sample multiplexing for retinal single-cell RNA-sequencing"

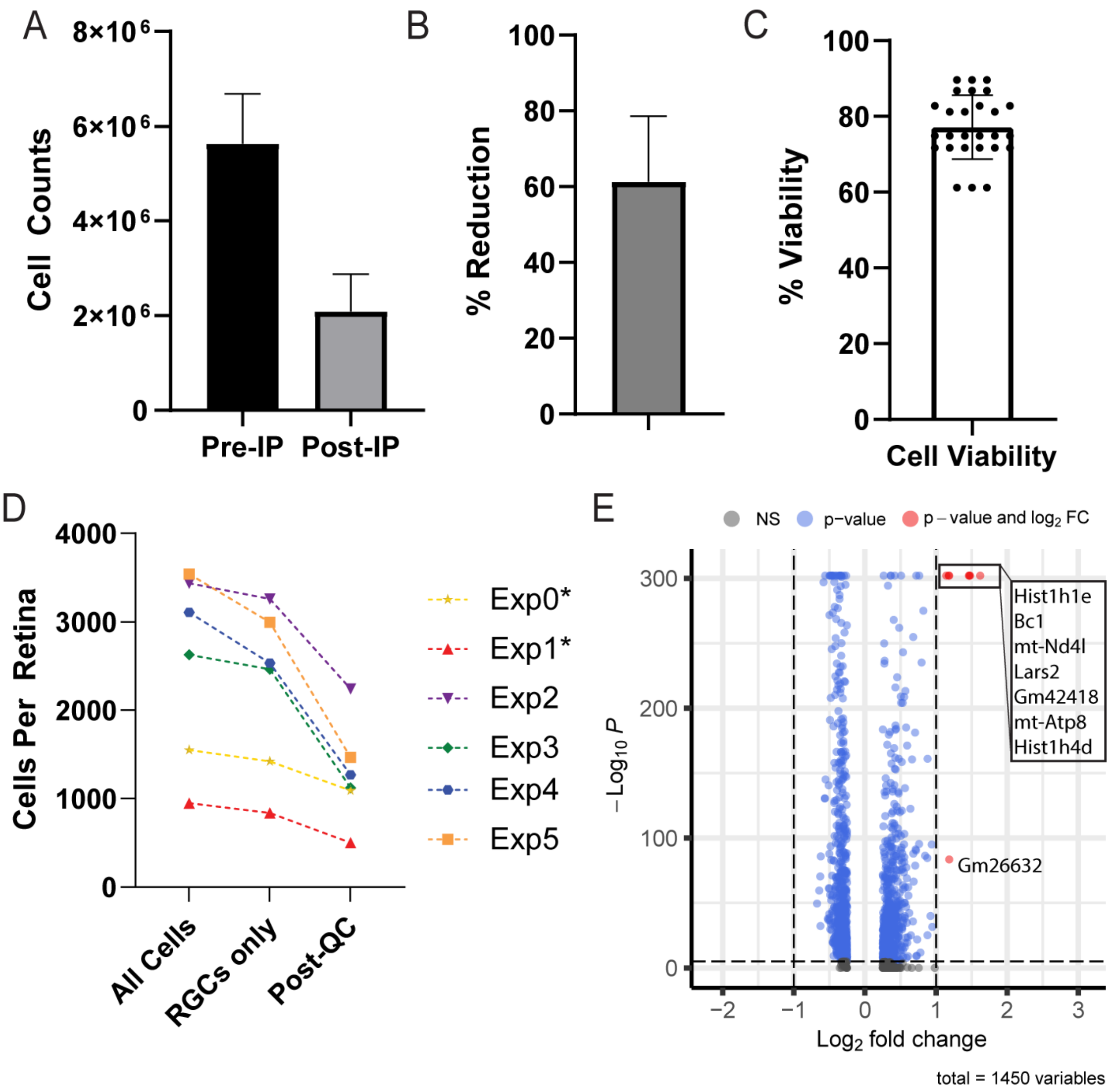


<u>**Figure S1. Evaluation of the CMO-Multiplexing Protocol.**</u> (A-B) Representation of the amount of cells removed by negative-rod immunopanning. Cells were reduced from ~6 x $10^6$ to ~2 x $10^6$, which is approximately a 60% reduction. n = 27 retinas. (C) Representation of the cell viability for all processed retinas, as determined by DAPI staining during FACS. The average viability was ~77± 8.3%. (D) Depiction of RGC recovery per experiment. Discounting Exp0 and Exp1 as non-representative data (only a portion of cells were sequenced), the average RGC recovery was 1466 cells per retina. (E) Comparison of DEG analysis between Unlabeled and CMO-labeled data. (Relevant to Figure 1).

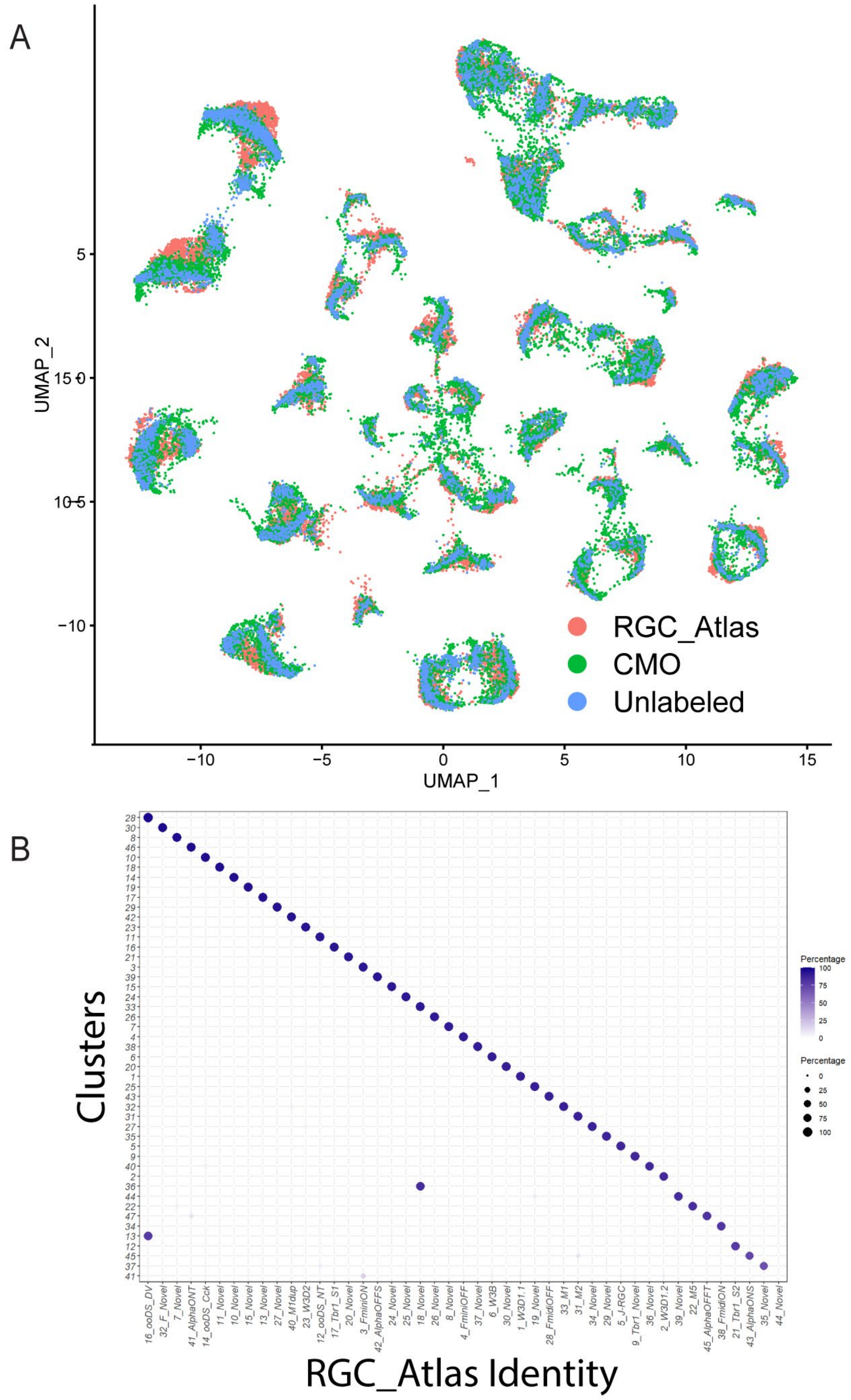


**<u>Figure S2. Evaluation of Transcriptomic Correspondence to the RGC Atlas.</u>** (A) UMAP of an integrated dataset between the RGC_Atlas, CMO-labeled data, and Unlabeled data. (B) Confusion matrix comparing refined clusters from the CMO dataset to the RGC Atlas clusters. Data indicates high transcriptomic cluster to cluster correspondence except for 44_Novel.

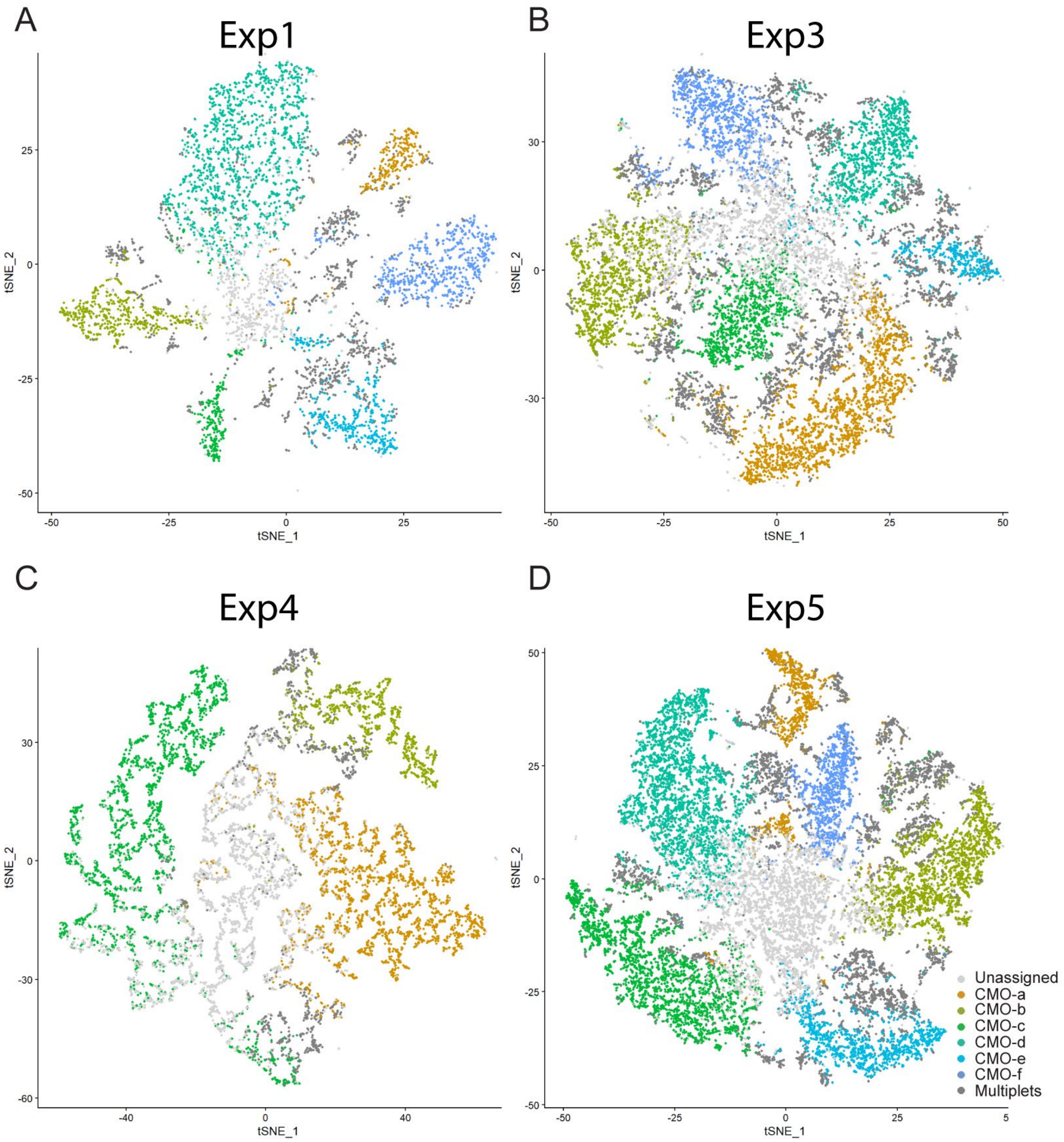


**Figure S3. CMO-based Clustering and Assignments.** (A-D) tSNE showing clustering of cells from Experiments 1, 3, 4, and 5 based on CMO-barcode reads. (Relevant to Figure 2).

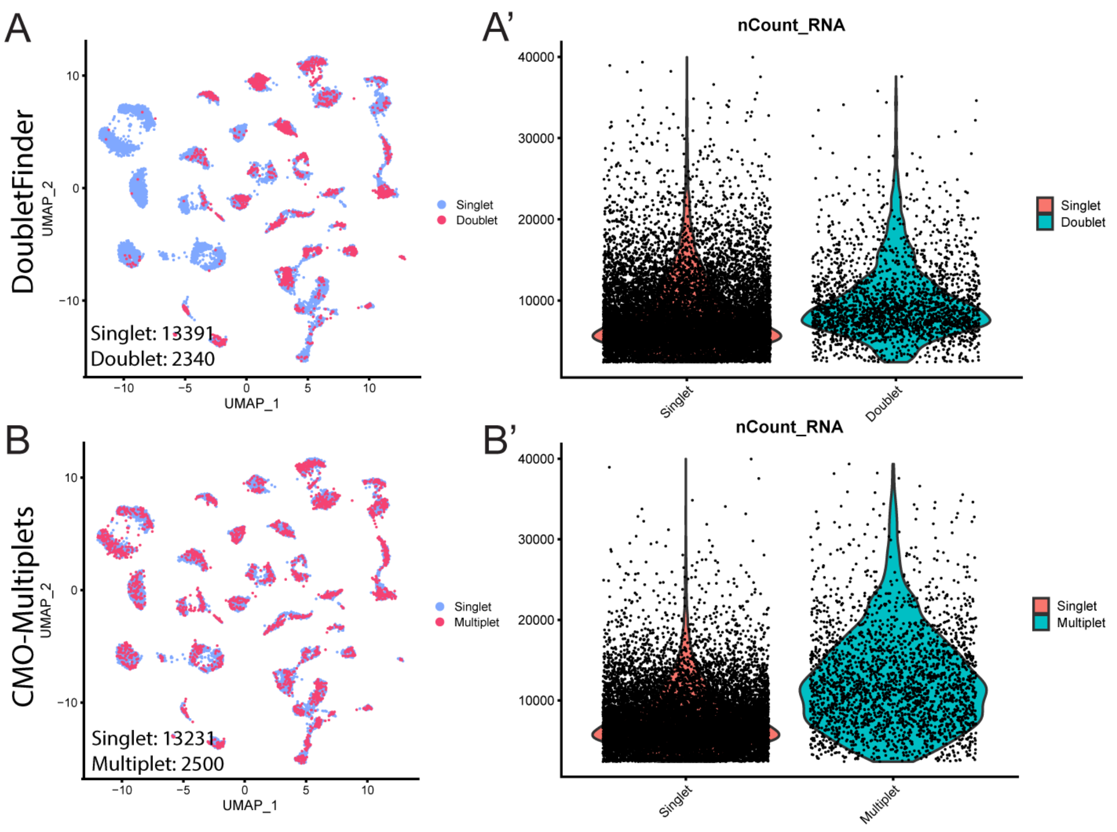


**<u>Figure S4. Comparison of Multiplet Identification of DoubletFinder versus CMO-defined Multiplets.</u>** Comparison of CMO-multiplets vs DoubletFinder doublets was performed on representative data from Experiment 2. (A-A') UMAP representation of doublets identified using DoubletFinder and a comparison of singlet versus doublet cell counts. (B-B') UMAP representation of multiplets identified using CMOs and a comparison of singlet versus multiplet cell counts. A similar number of multiplets are identified by both methods. DoubletFinder multiplet populations are highly localized in a few clusters, while CMO-defined multiplets are evenly distributed. The range of counts identified by CMOs is also wider, suggesting proper detection of multiplets, rather than only doublets.

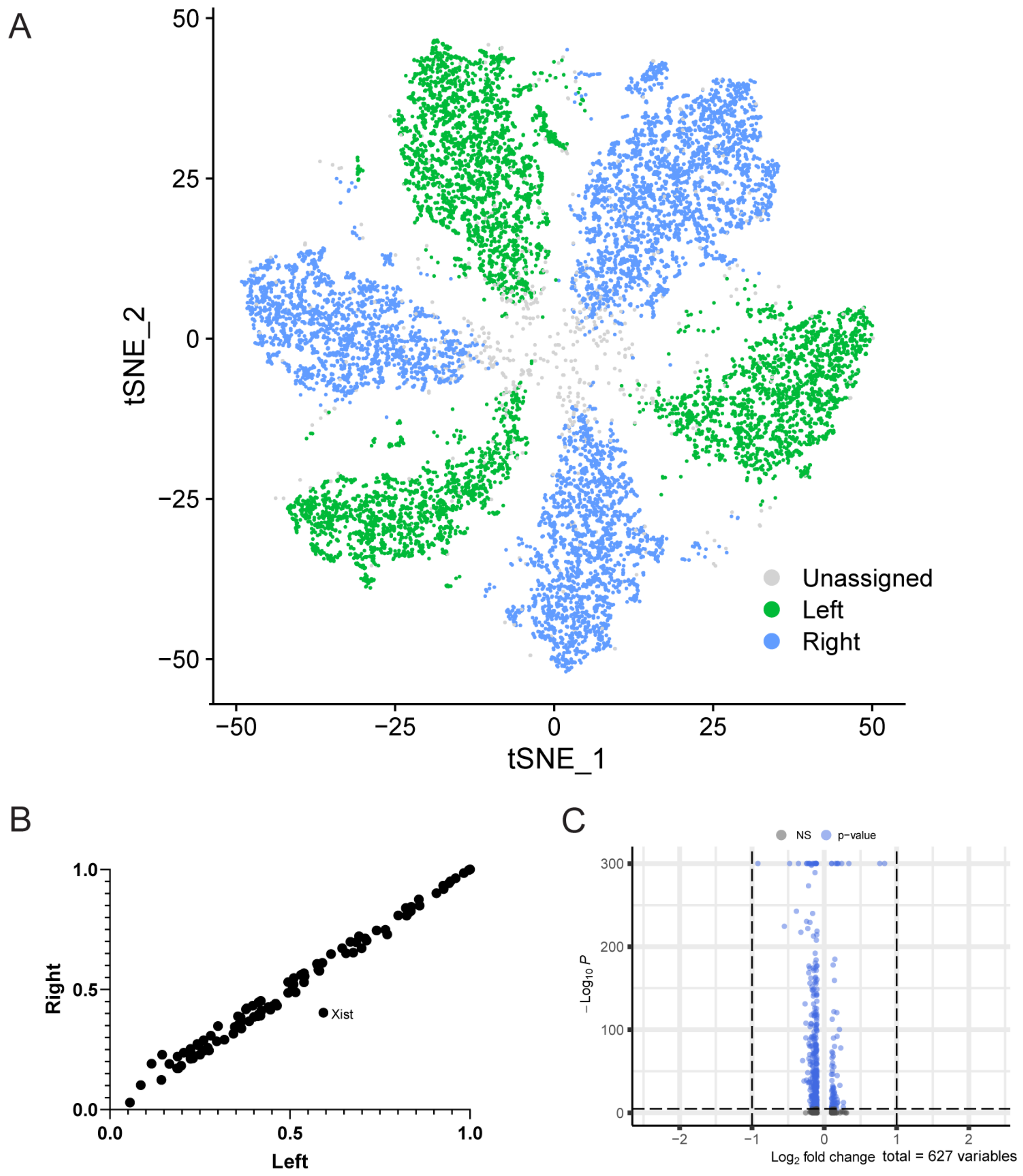


**Figure S5. Transcriptomic Differences are Minimal between Retinas derived from Left and Right Eyes.** (A) tSNE CMO plot showing distribution of left versus right eye derived RGCs based on CMO classification in Experiment 2. (B) Top DEG analysis comparing RGCs derived from left and right retinas plotted by percentage of cells with detected transcripts. (C) Volcano plot of top DEGs (FC: fold change, NS: not significant). No DEGs were found.

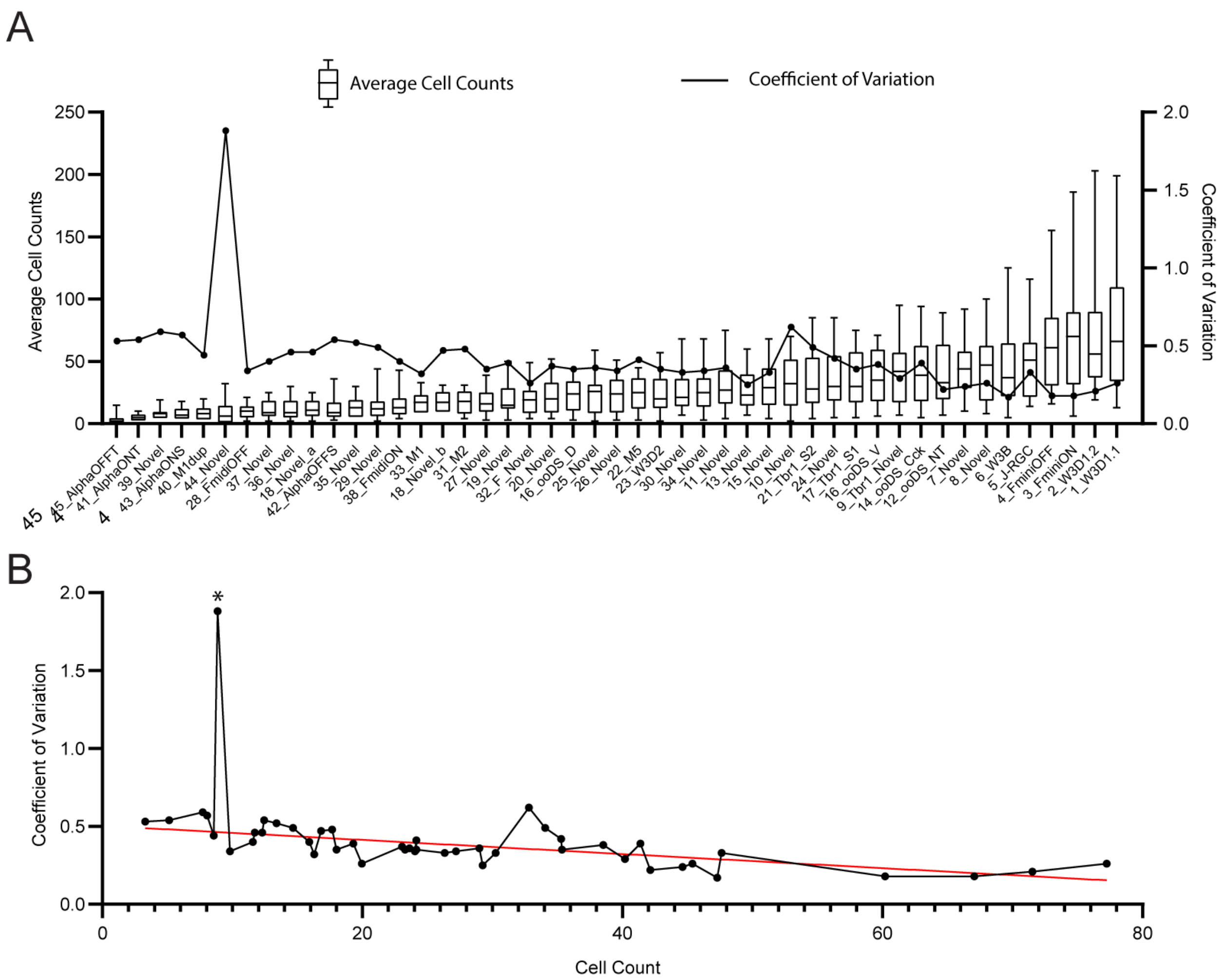


**<u>Figure S6. Cluster-Specific Marker Gene Variance Negatively Correlates with Cluster Size.</u>** (A) Integrated line and box plot depicting average cell counts (box) and marker gene variance per cluster between retinas (line). (21 retinas, Exp1 not included). Gene variance is portrayed using CV calculated using the average gene expression of the cluster-specific markers per cluster of a retina. (B) Scatter plot of Cell Counts vs Coefficient of Variation. The trendline indicates a trend of decreasing variance with increasing cell counts. *Cluster 44_Novel which was not well-represented by markers was not factored into the calculations of the trendline. (Relevant to Figure 5).

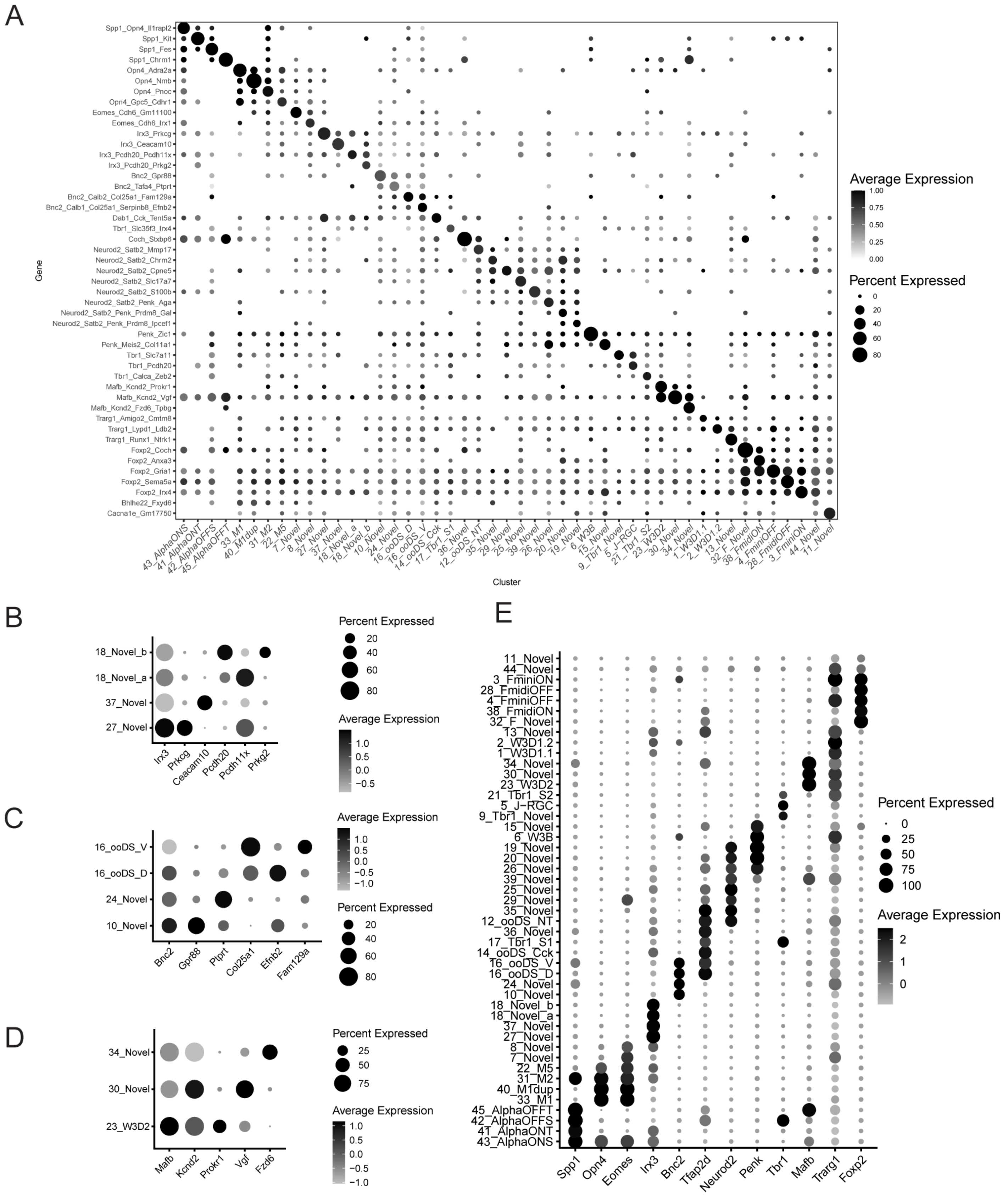

A
Average Expression
1.00
0.75
0.50
0.25
0.00
Percent Expressed
0
20
40
60
80
Gene
Cluster
Spp1_Opn4_Il1rapl2
Spp1_Kit
Spp1_Fes
Spp1_Chrm1
Opn4_Adra2a
Opn4_Nmb
Opn4_Pnoc
Opn4_Gpc5_Cdhr1
Eomes_Cdh6_Gm11100
Eomes_Cdh6_Irx1
Irx3_Prkcg
Irx3_Ceacam10
Irx3_Pcdh20_Pcdh11x
Irx3_Pcdh20_Prkg2
Bnc2_Gpr88
Bnc2_Tafa4_Ptprt
Bnc2_Calb2_Col25a1_Fam129a
Bnc2_Calb1_Col25a1_Serpinb8_Efnb2
Dab1_Cck_Tent5a
Tbr1_Slc35f3_Irx4
Coch_Stxbp6
Neurod2_Satb2_Mmp17
Neurod2_Satb2_Chrm2
Neurod2_Satb2_Cpne5
Neurod2_Satb2_Slc17a7
Neurod2_Satb2_S100b
Neurod2_Satb2_Penk_Aga
Neurod2_Satb2_Penk_Prdm8_Gal
Neurod2_Satb2_Penk_Prdm8_Ipcef1
Penk_Zic1
Penk_Meis2_Col11a1
Tbr1_Slc7a11
Tbr1_Pcdh20
Tbr1_Calca_Zeb2
Mafb_Kcnd2_Prokr1
Mafb_Kcnd2_Vgf
Mafb_Kcnd2_Fzd6_Tpbg
Trarg1_Amigo2_Cmtm8
Trarg1_Lypd1_Ldb2
Trarg1_Runx1_Ntrk1
Foxp2_Coch
Foxp2_Anxa3
Foxp2_Gria1
Foxp2_Sema5a
Foxp2_Irx4
Bhlhe22_Fxyd6
Cacna1e_Gm17750
43_AlphaONS
41_AlphaONT
42_AlphaOFFS
45_AlphaOFFT
33_M1
40_M1dup
31_M2
22_M5
7_Novel
8_Novel
27_Novel
37_Novel
18_Novel_a
18_Novel_b
10_Novel
24_Novel
16_ooDS_D
16_ooDS_V
14_ooDS_Cck
17_Tbr1_S1
36_Novel
12_ooDS_NT
35_Novel
29_Novel
25_Novel
39_Novel
26_Novel
20_Novel
19_Novel
6_W3B
15_Novel
9_Tbr1_Novel
5_J-RGC
21_Tbr1_S2
23_W3D2
30_Novel
34_Novel
1_W3D1.1
2_W3D1.2
13_Novel
32_F_Novel
38_FmidiON
4_FminiOFF
28_FmidiOFF
3_FminiON
44_Novel
11_Novel
B
18_Novel_b
18_Novel_a
37_Novel
27_Novel
Irx3
Prkcg
Ceacam10
Pcdh20
Pcdh11x
Prkg2
Percent Expressed
20
40
60
80
Average Expression
1.0
0.5
0.0
−0.5
C
16_ooDS_V
16_ooDS_D
24_Novel
10_Novel
Bnc2
Gpr88
Ptprt
Col25a1
Efnb2
Fam129a
Average Expression
1.0
0.5
0.0
−0.5
−1.0
Percent Expressed
20
40
60
80
D
34_Novel
30_Novel
23_W3D2
Mafb
Kcnd2
Prokr1
Vgf
Fzd6
Percent Expressed
25
50
75
Average Expression
1.0
0.5
0.0
−0.5
−1.0
E
11_Novel
44_Novel
3_FminiON
28_FmidiOFF
4_FminiOFF
38_FmidiON
32_F_Novel
13_Novel
2_W3D1.2
1_W3D1.1
34_Novel
30_Novel
23_W3D2
21_Tbr1_S2
5_J-RGC
9_Tbr1_Novel
15_Novel
6_W3B
19_Novel
20_Novel
26_Novel
39_Novel
25_Novel
29_Novel
35_Novel
12_ooDS_NT
36_Novel
17_Tbr1_S1
14_ooDS_Cck
16_ooDS_V
16_ooDS_D
24_Novel
10_Novel
18_Novel_b
18_Novel_a
37_Novel
27_Novel
8_Novel
7_Novel
22_M5
31_M2
40_M1dup
33_M1
45_AlphaOFFT
42_AlphaOFFS
41_AlphaONT
43_AlphaONS
Spp1
Opn4
Eomes
Irx3
Bnc2
Tfap2d
Neurod2
Penk
Tbr1
Mafb
Trarg1
Foxp2
Percent Expressed
0
25
50
75
100
Average Expression
2
1
0

**Figure S7. Specificity of comprehensive cluster-specific gene markers.** (A) Dotplot depicting gene marker combinations (rows), that are unique to each cluster (columns). Size represents percentage expression and shade represents expression level. Both Percentage Expression and Expression Level are averaged for the marker gene combinations. (B-D) Dotplots depicting specificity of markers for clusters characterized by their expression of specific gene groups: (B) *Irx3* (C) *Bnc2* (D) *Mafb/Kcnd2*. (E) Dotplot depicting markers expressed by RGC subgroups. (Relevant to Figure 6, Table 1).

**Table S1. Overview of RGC recovery per retina** (relevant to Figure 1)

| Experiment # | Cells Recovered post-Sequencing | # of Retinas | Average Cells Per Retina | RGCs Recovered | RGC Purity | Average Viability | Average RGCs Per Retina | RGCs Retained Post-QC | Post-QC per Retina |
|---|---|---|---|---|---|---|---|---|---|
| Exp0_Unlabeled* | 12405 | 8 | 1551 | 11377 | 92% | 88% | 1422 | 8734 | 1092 |
| Exp1_CMO* | 5703 | 6 | 951 | 5031 | 88% | 72% | 839 | 3016 | 503 |
| Exp2_CMO | 20623 | 6 | 3437 | 19552 | 95% | 75% | 3259 | 13311 | 2219 |
| Exp3_CMO | 15750 | 6 | 2625 | 14777 | 94% | 74% | 2463 | 6236 | 1039 |
| Exp4_CMO | 9318 | 3 | 3106 | 7602 | 82% | 90% | 2534 | 3826 | 1275 |
| Exp5_CMO | 21246 | 6 | 3541 | 17979 | 85% | 82% | 2997 | 7985 | 1331 |

*Outliers: Exp0 and Exp1 had reduced input of cell numbers for sequencing.

**Table S2. Overview of RGC Recovery per CMO** (relevant to Figure 2)

Pre-QC

| Experiment # | Unassigned | Multiplet | CMO-a | CMO-b | CMO-c | CMO-d | CMO-e | CMO-f | Total Cells |
|---|---|---|---|---|---|---|---|---|---|
| Exp0_Unlabeled | n/a | n/a | n/a | n/a | n/a | n/a | n/a | n/a | 12405 |
| Exp1_CMO | 778 | 1361 | 298 | 490 | 270 | 1352 | 467 | 687 | 5703 |
| Exp2_CMO | 1955 | 3078 | 2204 | 3375 | 2922 | 2352 | 2331 | 2406 | 20623 |
| Exp3_CMO | 3254 | 4339 | 2354 | 1451 | 1278 | 1271 | 519 | 1284 | 15750 |
| Exp4_CMO | 3273 | 1019 | 2069 | 847 | 2110 | 0 | 0 | 0 | 9318 |
| Exp5_CMO | 4919 | 4567 | 990 | 2196 | 2766 | 3111 | 1417 | 1280 | 21246 |

Post-QC

| Experiment # | Unassigned | Multiplet | CMO-a | CMO-b | CMO-c | CMO-d | CMO-e | CMO-f | Total Cells |
|---|---|---|---|---|---|---|---|---|---|
| Exp0_Unlabeled | n/a | n/a | n/a | n/a | n/a | n/a | n/a | n/a | 8734 |
| Exp1_CMO | 291 | 0 | 181 | 337 | 195 | 1163 | 303 | 546 | 3016 |
| Exp2_CMO | 696 | 0 | 1702 | 2863 | 2494 | 1885 | 1762 | 1909 | 13311 |
| Exp3_CMO | 1233 | 0 | 1482 | 917 | 850 | 729 | 267 | 758 | 6236 |
| Exp4_CMO | 1032 | 0 | 1190 | 388 | 1216 | 0 | 0 | 0 | 3826 |
| Exp5_CMO | 1175 | 0 | 431 | 1398 | 1805 | 2022 | 648 | 506 | 7985 |

**Table S3. Percent Expression of Major Sex-Related DEGs** (relevant to Figure 3)

| | CMO | Xist | Ddx3y | Eif2s3y | Label | Counts | % Counts per Exp |
|---|---|---|---|---|---|---|---|
| Exp0 | nonCMO | 72.2% | 12.0% | 16.3% | n/a | 8734 | 100% |
| Exp1* | CMO-a | 99.4% | 0.0% | 0.6% | F | 181 | 6% |
| | CMO-b | 99.4% | 0.0% | 1.5% | F | 337 | 11% |
| | CMO-c | 99.5% | 0.0% | 1.5% | F | 195 | 6% |
| | CMO-d | 99.7% | 0.8% | 0.5% | F | 1163 | 39% |
| | CMO-e | 9.9% | 51.5% | 69.0% | M | 303 | 10% |
| | CMO-f | 5.9% | 50.4% | 69.0% | M | 546 | 18% |
| | Negative | 88.7% | 6.9% | 8.6% | n/a | 291 | 10% |
| Exp2 | CMO-a | 5.3% | 27.1% | 34.3% | M | 1702 | 13% |
| | CMO-b | 3.9% | 31.2% | 37.3% | M | 2863 | 22% |
| | CMO-c | 96.1% | 1.0% | 1.0% | F | 2494 | 19% |
| | CMO-d | 95.8% | 0.7% | 1.0% | F | 1885 | 14% |
| | CMO-e | 95.6% | 1.0% | 1.0% | F | 1762 | 13% |
| | CMO-f | 95.1% | 1.0% | 1.2% | F | 1909 | 14% |
| | Negative | 64.7% | 9.9% | 14.1% | n/a | 696 | 5% |
| Exp3 | CMO-a | 0.0% | 24.6% | 34.3% | M | 1482 | 24% |
| | CMO-b | 0.0% | 25.8% | 33.6% | M | 917 | 15% |
| | CMO-c | 0.0% | 26.8% | 34.8% | M | 850 | 14% |
| | CMO-d | 0.0% | 24.8% | 33.7% | M | 729 | 12% |
| | CMO-e** | 0.4% | 27.0% | 30.7% | M | 267 | 4% |
| | CMO-f | 0.0% | 26.8% | 33.6% | M | 758 | 12% |
| | Negative | 0.0% | 22.5% | 32.3% | n/a | 1233 | 32% |
| Exp4 | CMO-a | 99.6% | 2.6% | 0.3% | F | 1190 | 31% |
| | CMO-b** | 24.5% | 49.5% | 59.5% | M | 388 | 10% |
| | CMO-c | 99.8% | 3.1% | 0.5% | F | 1216 | 32% |
| | Negative | 96.8% | 2.6% | 1.6% | n/a | 1032 | 27% |
| Exp5 | CMO-a** | 24.6% | 24.8% | 38.5% | M | 431 | 5% |
| | CMO-b | 97.1% | 2.9% | 0.7% | F | 1398 | 18% |
| | CMO-c | 97.2% | 1.9% | 1.0% | F | 1805 | 23% |
| | CMO-d | 97.7% | 2.5% | 0.8% | F | 2022 | 25% |
| | CMO-e** | 18.1% | 31.3% | 37.0% | M | 648 | 8% |
| | CMO-f** | 13.4% | 31.6% | 47.4% | M | 506 | 6% |
| | Negative | 84.0% | 4.7% | 5.8% | n/a | 1175 | 15% |

* Exp1 has some cell losses from isolation procedure, but CMO-classification was unaffe

**Outliers: CMO-labeling efficiencies were lower for these samples, which may have impacted assignment accuracy.

**Table S4. RGC Type Representation** (relevant to Figure 4)

| | | | **CMO_Assigned Statistics*** | | | | | | |
|---|---|---|---|---|---|---|---|---|---|
| Cluster | RGC_Atlas | Exp0_Unlabeled | Unassigned | Assigned | Average | Std Dev | Median | Min | Max |
| 43_AlphaONS | 106 | 23 | 30 | 179 | 8.05 | 5.04 | 7 | 0 | 18 |
| 41_AlphaONT | 126 | 18 | 24 | 114 | 5.14 | 2.71 | 5 | 1 | 10 |
| 42_AlphaOFFS | 113 | 24 | 67 | 276 | 12.43 | 8.72 | 9 | 3 | 36 |
| 45_AlphaOFFT | 54 | 4 | 15 | 69 | 3.29 | 3.51 | 2 | 0 | 15 |
| 33_M1 | 323 | 144 | 49 | 396 | 16.29 | 8.84 | 17 | 0 | 33 |
| 40_M1dup | 174 | 70 | 38 | 206 | 8.57 | 4.98 | 8 | 1 | 20 |
| 31_M2 | 444 | 149 | 51 | 414 | 17.67 | 9.24 | 18 | 4 | 31 |
| 22_M5 | 610 | 176 | 86 | 553 | 24.14 | 13.26 | 25 | 3 | 45 |
| 7_Novel | 1579 | 359 | 201 | 1039 | 44.62 | 21.83 | 44 | 10 | 92 |
| 8_Novel | 1258 | 353 | 200 | 1055 | 45.38 | 25.39 | 47 | 8 | 100 |
| 27_Novel | 529 | 169 | 86 | 417 | 18.00 | 10.47 | 16 | 3 | 39 |
| 37_Novel | 213 | 86 | 69 | 266 | 11.57 | 6.61 | 9 | 2 | 25 |
| 18_Novel_a | 423** | 107 | 62 | 284 | 12.29 | 7.34 | 11 | 2 | 25 |
| 18_Novel_b | 423** | 109 | 65 | 379 | 16.81 | 9.01 | 17 | 0 | 31 |
| 10_Novel | 1170 | 253 | 165 | 763 | 32.81 | 20.37 | 32 | 2 | 70 |
| 24_Novel | 553 | 185 | 160 | 788 | 35.29 | 22.30 | 30 | 5 | 85 |
| 16_ooDS_D | 415** | 102 | 86 | 521 | 23.29 | 12.47 | 24 | 3 | 44 |
| 16_ooDS_V | 414** | 159 | 146 | 853 | 38.52 | 21.60 | 35 | 6 | 71 |
| 14_ooDS_Cck | 875 | 289 | 131 | 935 | 41.38 | 25.59 | 39 | 5 | 94 |
| 17_Tbr1_S1 | 828 | 141 | 200 | 785 | 35.33 | 22.05 | 30 | 5 | 75 |
| 36_Novel | 236 | 52 | 37 | 265 | 11.71 | 8.26 | 9 | 2 | 30 |
| 12_ooDS_NT | 953 | 212 | 96 | 948 | 42.14 | 25.03 | 33 | 7 | 89 |
| 35_Novel | 310 | 96 | 31 | 320 | 13.38 | 7.52 | 13 | 1 | 30 |
| 29_Novel | 499 | 92 | 62 | 337 | 14.67 | 11.89 | 12 | 2 | 44 |
| 25_Novel | 542 | 106 | 75 | 529 | 23.62 | 14.33 | 26 | 2 | 59 |
| 39_Novel | 202 | 37 | 43 | 178 | 7.71 | 4.29 | 8 | 1 | 19 |
| 26_Novel | 534 | 143 | 50 | 538 | 24.05 | 15.31 | 24 | 3 | 51 |
| 20_Novel | 711 | 195 | 125 | 545 | 23.05 | 14.58 | 20 | 4 | 52 |
| 19_Novel | 775 | 175 | 132 | 455 | 19.29 | 11.68 | 15 | 3 | 50 |
| 6_W3B | 1590 | 368 | 299 | 1096 | 47.29 | 31.91 | 37 | 5 | 125 |
| 15_Novel | 865 | 144 | 102 | 690 | 30.24 | 17.49 | 29 | 4 | 68 |
| 9_Tbr1_Novel | 1223 | 381 | 137 | 965 | 40.19 | 24.72 | 42 | 7 | 95 |
| 5_J-RGC | 1715 | 323 | 249 | 1093 | 47.62 | 26.42 | 51 | 14 | 116 |
| 21_Tbr1_S2 | 687 | 230 | 268 | 780 | 34.05 | 21.56 | 28 | 4 | 85 |
| 23_W3D2 | 601 | 100 | 180 | 544 | 24.14 | 14.22 | 20 | 2 | 57 |
| 30_Novel | 491 | 118 | 188 | 593 | 26.33 | 15.51 | 21 | 7 | 68 |
| 34_Novel | 312 | 76 | 114 | 605 | 27.19 | 17.03 | 25 | 3 | 68 |
| 1_W3D1.1 | 3000 | 890 | 412 | 1942 | 77.24 | 51.33 | 66 | 13 | 199 |
| 2_W3D1.2 | 2859 | 413 | 670 | 1662 | 71.52 | 45.67 | 56 | 19 | 203 |
| 13_Novel | 943 | 269 | 173 | 675 | 29.24 | 16.63 | 23 | 7 | 60 |
| 32_F_Novel | 407 | 75 | 50 | 442 | 19.95 | 13.00 | 19 | 4 | 49 |
| 38_FmidiON | 207 | 69 | 79 | 361 | 15.90 | 10.44 | 13 | 4 | 43 |
| 4_FminiOFF | 1868 | 404 | 357 | 1432 | 60.19 | 35.41 | 61 | 16 | 155 |
| 28_FmidiOFF | 517 | 60 | 23 | 222 | 9.81 | 5.33 | 10 | 2 | 21 |
| 3_FminiON | 1990 | 585 | 411 | 1589 | 67.05 | 44.70 | 70 | 6 | 186 |
| 44_Novel | 62 | 7 | 33 | 188 | 8.86 | 9.69 | 6 | 0 | 32 |
| 11_Novel | 990 | 194 | 151 | 661 | 29.00 | 16.74 | 27 | 4 | 75 |
| SUM | 34044 | 8734 | 6478 | 29947 | 1296 | 792 | 1189 | 209 | 3088 |

*Exp1 is excluded because of a low cell input for sequencing.

**The RGC_Atlas does not differentiate these neighboring clusters so the values assigned to each is half of the total.

**Table S5. High and Low Variance Genes Between Retinas** (relevant to Figure 5)

| High Variance | Mean* | StdDev | CV | Low Variance | Mean* | StdDev | CV |
|---|---|---|---|---|---|---|---|
| Tsix | 0.10 | 0.12 | 125.68 | Slc38a1 | 1.96 | 0.09 | 4.64 |
| Eif2s3y | 0.25 | 0.31 | 124.07 | Fra10ac1 | 0.55 | 0.03 | 4.62 |
| Tac2 | 0.07 | 0.09 | 122.88 | Zc3h15 | 3.32 | 0.15 | 4.58 |
| Gad1 | 0.09 | 0.11 | 116.91 | Tmem167 | 1.15 | 0.05 | 4.58 |
| Uty | 0.09 | 0.11 | 116.11 | Ube2b | 3.08 | 0.14 | 4.55 |
| Xist | 3.75 | 4.19 | 111.71 | Matr3 | 2.15 | 0.10 | 4.55 |
| Kdm5d | 0.06 | 0.07 | 111.50 | Elavl2 | 2.11 | 0.10 | 4.52 |
| Ddx3y | 0.15 | 0.16 | 110.02 | Kif1a | 3.16 | 0.14 | 4.47 |
| Slc32a1 | 0.06 | 0.06 | 105.73 | Atp1b1 | 13.68 | 0.61 | 4.46 |
| C1ql2 | 0.05 | 0.06 | 104.82 | Pls3 | 1.33 | 0.06 | 4.45 |
| Tfap2b | 0.07 | 0.07 | 101.31 | Cul5 | 0.96 | 0.04 | 4.40 |
| Gldn | 0.06 | 0.05 | 88.32 | Kif5b | 2.59 | 0.11 | 4.37 |
| Cplx3 | 0.20 | 0.16 | 78.30 | Cdc42 | 4.36 | 0.19 | 4.33 |
| Nppb | 0.09 | 0.07 | 77.63 | Tpm3 | 1.49 | 0.06 | 4.30 |
| Xlr3b | 0.06 | 0.04 | 71.90 | Thoc7 | 2.13 | 0.09 | 4.29 |
| Cdkn1c | 0.16 | 0.11 | 71.73 | Napg | 1.68 | 0.07 | 4.25 |
| Lpcat4 | 0.09 | 0.07 | 71.60 | Hsp90aa1 | 16.93 | 0.71 | 4.19 |
| Hist1h4d | 0.13 | 0.10 | 71.43 | Impa1 | 1.21 | 0.05 | 4.16 |
| C1ql1 | 0.17 | 0.11 | 64.40 | Eif2s2 | 3.16 | 0.13 | 4.15 |
| Rho | 0.11 | 0.07 | 62.47 | Sdf4 | 0.80 | 0.03 | 4.10 |
| Gm21887 | 0.08 | 0.05 | 62.33 | Pja2 | 3.26 | 0.13 | 3.92 |
| Gm42418 | 33.36 | 20.67 | 61.98 | Yaf2 | 1.93 | 0.08 | 3.90 |
| Lbhd2 | 0.05 | 0.03 | 60.83 | Rsrc2 | 1.91 | 0.07 | 3.82 |
| A430106G13Rik | 0.05 | 0.03 | 60.70 | Thra | 4.02 | 0.15 | 3.71 |
| Pcdhga12 | 0.08 | 0.05 | 60.54 | Chn1 | 3.29 | 0.11 | 3.45 |
| Cmss1 | 0.28 | 0.17 | 58.45 | Ttc3 | 20.22 | 0.70 | 3.44 |
| Gm17167 | 0.08 | 0.05 | 58.36 | Pura | 4.40 | 0.15 | 3.30 |
| Hist1h1e | 0.27 | 0.16 | 56.58 | Usp14 | 2.04 | 0.07 | 3.29 |
| Frs3 | 0.09 | 0.05 | 56.39 | Pax6 | 3.04 | 0.09 | 2.87 |
| Lars2 | 1.34 | 0.72 | 54.21 | Srpk2 | 3.38 | 0.09 | 2.75 |

*only includes genes above scaled-expression mean value of 0.05.

**Table S6. Statistics for Cluster-Specific Marker Gene Variance** (relevant to Figure S6)

| Order | Cluster | Marker Gene | min | max | med | mean | SD | CV | Order | Cluster | Marker Gene | min | max | med | mean | SD | CV |
|---|---|---|---|---|---|---|---|---|---|---|---|---|---|---|---|---|---|
| 1 | 43_AlphaONS | Spp1 | 3.21 | 41.22 | 14.92 | 15.41 | 8.29 | 0.54 | 66 | 25_Novel | Neurod2 | 1.15 | 2.74 | 1.50 | 1.60 | 0.40 | 0.25 |
| 2 | 43_AlphaONS | Opn4 | 0.44 | 2.01 | 1.09 | 1.11 | 0.43 | 0.39 | 67 | 25_Novel | Satb2 | 0.26 | 1.92 | 1.05 | 1.10 | 0.37 | 0.34 |
| 3 | 43_AlphaONS | Il1rapl2 | 0.00 | 0.91 | 0.22 | 0.31 | 0.23 | 0.73 | 68 | 25_Novel | Slc17a7 | 0.00 | 1.67 | 0.72 | 0.87 | 0.41 | 0.47 |
| 4 | 41_AlphaONT | Spp1 | 1.98 | 20.26 | 8.55 | 8.37 | 4.22 | 0.50 | 69 | 39_Novel | Neurod2 | 0.20 | 2.70 | 0.65 | 0.76 | 0.57 | 0.75 |
| 5 | 41_AlphaONT | Kit | 0.00 | 2.53 | 1.11 | 1.16 | 0.65 | 0.56 | 70 | 39_Novel | Satb2 | 0.00 | 1.28 | 0.49 | 0.50 | 0.29 | 0.58 |
| 6 | 42_AlphaOFFS | Spp1 | 8.07 | 33.02 | 16.33 | 17.95 | 7.32 | 0.41 | 71 | 39_Novel | S100b | 0.35 | 3.75 | 2.29 | 2.29 | 0.90 | 0.39 |
| 7 | 42_AlphaOFFS | Fes | 0.08 | 1.32 | 0.37 | 0.47 | 0.31 | 0.65 | 72 | 26_Novel | Neurod2 | 0.24 | 1.01 | 0.58 | 0.62 | 0.20 | 0.33 |
| 8 | 45_AlphaOFFT | Spp1 | 3.82 | 35.08 | 16.39 | 17.17 | 8.42 | 0.49 | 73 | 26_Novel | Satb2 | 0.72 | 1.75 | 1.17 | 1.15 | 0.27 | 0.23 |
| 9 | 45_AlphaOFFT | Chrm1 | 0.00 | 1.55 | 0.81 | 0.77 | 0.42 | 0.55 | 74 | 26_Novel | Penk | 0.70 | 4.13 | 1.76 | 1.91 | 0.78 | 0.41 |
| 10 | 33_M1 | Opn4 | 4.22 | 9.30 | 5.59 | 5.75 | 1.28 | 0.22 | 75 | 26_Novel | Aga | 0.50 | 2.26 | 0.98 | 1.06 | 0.39 | 0.37 |
| 11 | 33_M1 | Adra2a | 0.21 | 2.69 | 1.33 | 1.29 | 0.52 | 0.40 | 76 | 20_Novel | Neurod2 | 0.29 | 1.77 | 1.01 | 1.06 | 0.38 | 0.36 |
| 12 | 40_M1dup | Opn4 | 4.95 | 18.37 | 10.00 | 10.19 | 3.08 | 0.30 | 77 | 20_Novel | Satb2 | 0.52 | 2.09 | 1.13 | 1.16 | 0.38 | 0.33 |
| 13 | 40_M1dup | Nmb | 0.67 | 7.12 | 2.11 | 2.34 | 1.31 | 0.56 | 78 | 20_Novel | Penk | 2.16 | 6.26 | 3.51 | 3.64 | 0.96 | 0.26 |
| 14 | 31_M2 | Opn4 | 1.65 | 3.88 | 2.85 | 2.82 | 0.67 | 0.24 | 79 | 20_Novel | Prdm8 | 1.14 | 4.66 | 2.85 | 2.87 | 0.89 | 0.31 |
| 15 | 31_M2 | Pnoc | 0.00 | 0.74 | 0.26 | 0.33 | 0.23 | 0.70 | 80 | 20_Novel | Gal | 0.21 | 2.50 | 0.83 | 0.89 | 0.50 | 0.56 |
| 16 | 22_M5 | Opn4 | 0.00 | 1.57 | 0.91 | 0.88 | 0.39 | 0.45 | 81 | 19_Novel | Neurod2 | 0.41 | 2.07 | 1.07 | 1.08 | 0.37 | 0.35 |
| 17 | 22_M5 | Gpc5 | 0.00 | 1.40 | 0.90 | 0.85 | 0.29 | 0.34 | 82 | 19_Novel | Satb2 | 0.22 | 1.59 | 0.69 | 0.75 | 0.33 | 0.44 |
| 18 | 22_M5 | Cdhr1 | 0.27 | 1.43 | 0.77 | 0.76 | 0.31 | 0.41 | 83 | 19_Novel | Penk | 1.34 | 6.28 | 3.59 | 3.73 | 1.04 | 0.28 |
| 19 | 7_Novel | Eomes | 0.71 | 2.08 | 1.33 | 1.26 | 0.31 | 0.24 | 84 | 19_Novel | Prdm8 | 2.19 | 5.46 | 3.01 | 3.34 | 0.95 | 0.28 |
| 20 | 7_Novel | Cdh6 | 1.22 | 2.77 | 2.29 | 2.26 | 0.37 | 0.17 | 85 | 19_Novel | Ipcef1 | 0.00 | 0.61 | 0.27 | 0.25 | 0.14 | 0.56 |
| 21 | 7_Novel | Gm11100 | 0.44 | 1.83 | 1.13 | 1.07 | 0.31 | 0.29 | 86 | 6_W3B | Penk | 2.15 | 4.79 | 3.17 | 3.14 | 0.62 | 0.20 |
| 22 | 8_Novel | Eomes | 0.33 | 1.19 | 0.69 | 0.72 | 0.18 | 0.24 | 87 | 6_W3B | Zic1 | 2.45 | 3.90 | 3.16 | 3.15 | 0.40 | 0.13 |
| 23 | 8_Novel | Cdh6 | 0.39 | 1.66 | 1.24 | 1.19 | 0.30 | 0.25 | 88 | 15_Novel | Penk | 0.73 | 2.84 | 1.71 | 1.77 | 0.49 | 0.28 |
| 24 | 8_Novel | Irx1 | 0.38 | 1.03 | 0.61 | 0.61 | 0.15 | 0.25 | 89 | 15_Novel | Meis2 | 3.28 | 8.92 | 6.56 | 6.17 | 1.42 | 0.23 |
| 25 | 27_Novel | Irx3 | 1.15 | 2.59 | 1.69 | 1.71 | 0.38 | 0.22 | 90 | 15_Novel | Col11a1 | 0.00 | 1.20 | 0.55 | 0.55 | 0.25 | 0.46 |
| 26 | 27_Novel | Prkcg | 0.00 | 1.23 | 0.64 | 0.63 | 0.30 | 0.47 | 91 | 9_Tbr1_Novel | Tbr1 | 0.30 | 1.05 | 0.47 | 0.51 | 0.17 | 0.33 |
| 27 | 37_Novel | Irx3 | 0.27 | 2.00 | 1.06 | 1.10 | 0.35 | 0.32 | 92 | 9_Tbr1_Novel | Slc7a11 | 0.54 | 2.98 | 2.15 | 2.13 | 0.48 | 0.23 |
| 28 | 37_Novel | Ceacam10 | 0.19 | 0.96 | 0.57 | 0.55 | 0.25 | 0.45 | 93 | 5_J-RGC | Tbr1 | 0.28 | 1.04 | 0.65 | 0.69 | 0.22 | 0.32 |
| 29 | 18_Novel_a | Irx3 | 0.59 | 2.43 | 1.47 | 1.53 | 0.46 | 0.30 | 94 | 5_J-RGC | Pcdh20 | 0.29 | 1.01 | 0.58 | 0.58 | 0.18 | 0.32 |
| 30 | 18_Novel_a | Pcdh20 | 0.00 | 0.47 | 0.25 | 0.23 | 0.14 | 0.64 | 95 | 21_Tbr1_S2 | Tbr1 | 0.04 | 1.24 | 0.27 | 0.33 | 0.25 | 0.76 |
| 31 | 18_Novel_a | Pcdh11x | 0.62 | 3.55 | 1.73 | 1.82 | 0.74 | 0.41 | 96 | 21_Tbr1_S2 | Calca | 0.62 | 2.95 | 1.63 | 1.72 | 0.55 | 0.32 |
| 32 | 18_Novel_b | Irx3 | 0.68 | 1.91 | 1.22 | 1.23 | 0.32 | 0.26 | 97 | 21_Tbr1_S2 | Zeb2 | 0.54 | 2.90 | 1.36 | 1.41 | 0.51 | 0.36 |
| 33 | 18_Novel_b | Pcdh20 | 0.18 | 2.05 | 0.72 | 0.79 | 0.41 | 0.51 | 98 | 23_W3D2 | Mafb | 2.49 | 8.49 | 6.27 | 6.17 | 1.55 | 0.25 |
| 34 | 18_Novel_b | Prkg2 | 0.00 | 0.50 | 0.23 | 0.25 | 0.15 | 0.61 | 99 | 23_W3D2 | Kcnd2 | 2.57 | 6.79 | 4.12 | 4.37 | 1.28 | 0.29 |
| 35 | 10_Novel | Bnc2 | 0.14 | 2.19 | 0.40 | 0.48 | 0.40 | 0.84 | 100 | 23_W3D2 | Prokr1 | 0.11 | 1.53 | 0.66 | 0.68 | 0.32 | 0.47 |
| 36 | 10_Novel | Gpr88 | 0.31 | 1.78 | 0.95 | 0.93 | 0.35 | 0.37 | 101 | 30_Novel | Mafb | 1.80 | 5.30 | 3.42 | 3.58 | 0.99 | 0.28 |
| 37 | 24_Novel | Bnc2 | 0.00 | 0.53 | 0.34 | 0.34 | 0.12 | 0.35 | 102 | 30_Novel | Kcnd2 | 2.11 | 7.60 | 4.89 | 4.69 | 1.36 | 0.29 |
| 38 | 24_Novel | Tafa4 | 0.08 | 1.19 | 0.40 | 0.45 | 0.23 | 0.51 | 103 | 30_Novel | Vgf | 0.66 | 5.34 | 2.48 | 2.82 | 1.09 | 0.39 |
| 39 | 24_Novel | Ptprt | 0.18 | 1.17 | 0.81 | 0.75 | 0.27 | 0.36 | 104 | 34_Novel | Mafb | 0.60 | 5.61 | 3.75 | 3.70 | 1.00 | 0.27 |
| 40 | 16_ooDS_D | Bnc2 | 0.00 | 0.63 | 0.30 | 0.33 | 0.14 | 0.42 | 105 | 34_Novel | Kcnd2 | 1.62 | 5.51 | 3.80 | 3.38 | 1.14 | 0.34 |
| 41 | 16_ooDS_D | Calb2 | 32.33 | 63.17 | 45.27 | 46.27 | 8.41 | 0.18 | 106 | 34_Novel | Fzd6 | 0.30 | 1.14 | 0.57 | 0.66 | 0.25 | 0.38 |
| 42 | 16_ooDS_D | Col25a1 | 0.46 | 2.40 | 1.55 | 1.41 | 0.45 | 0.32 | 107 | 34_Novel | Tpbg | 0.31 | 1.71 | 0.95 | 0.93 | 0.32 | 0.35 |
| 43 | 16_ooDS_D | Fam129a | 0.09 | 1.10 | 0.49 | 0.50 | 0.23 | 0.46 | 108 | 1_W3D1.1 | Trarg1 | 2.54 | 4.96 | 3.17 | 3.25 | 0.55 | 0.17 |
| 44 | 16_ooDS_V | Bnc2 | 0.13 | 0.63 | 0.39 | 0.37 | 0.14 | 0.38 | 109 | 1_W3D1.1 | Amigo2 | 0.22 | 1.06 | 0.65 | 0.66 | 0.18 | 0.27 |
| 45 | 16_ooDS_V | Calb1 | 6.65 | 17.54 | 11.50 | 11.58 | 3.11 | 0.27 | 110 | 1_W3D1.1 | Cmtm8 | 0.18 | 0.79 | 0.38 | 0.42 | 0.14 | 0.33 |
| 46 | 16_ooDS_V | Col25a1 | 0.21 | 1.10 | 0.55 | 0.58 | 0.25 | 0.44 | 111 | 2_W3D1.2 | Trarg1 | 4.00 | 7.18 | 5.23 | 5.25 | 0.75 | 0.14 |
| 47 | 16_ooDS_V | Serpinb8 | 0.33 | 1.06 | 0.71 | 0.73 | 0.21 | 0.29 | 112 | 2_W3D1.2 | Lypd1 | 1.26 | 2.51 | 1.78 | 1.87 | 0.32 | 0.17 |
| 48 | 16_ooDS_V | Efnb2 | 0.33 | 2.18 | 0.88 | 1.02 | 0.46 | 0.46 | 113 | 2_W3D1.2 | Ldb2 | 0.21 | 0.84 | 0.43 | 0.47 | 0.14 | 0.30 |
| 49 | 14_ooDS_Cck | Dab1 | 0.00 | 1.65 | 0.84 | 0.86 | 0.35 | 0.41 | 114 | 13_Novel | Trarg1 | 1.77 | 4.10 | 2.36 | 2.45 | 0.53 | 0.22 |
| 50 | 14_ooDS_Cck | Cck | 0.62 | 4.77 | 3.01 | 3.16 | 0.94 | 0.30 | 115 | 13_Novel | Runx1 | 1.18 | 2.77 | 1.94 | 1.88 | 0.38 | 0.20 |
| 51 | 14_ooDS_Cck | Tent5a | 0.00 | 0.97 | 0.54 | 0.48 | 0.21 | 0.43 | 116 | 13_Novel | Ntrk1 | 0.37 | 1.73 | 1.15 | 1.13 | 0.35 | 0.31 |
| 52 | 17_Tbr1_S1 | Tbr1 | 0.51 | 1.50 | 0.84 | 0.91 | 0.25 | 0.27 | 117 | 32_F_Novel | Foxp2 | 2.27 | 5.86 | 3.04 | 3.23 | 0.79 | 0.24 |
| 53 | 17_Tbr1_S1 | Slc35f3 | 0.09 | 0.59 | 0.34 | 0.33 | 0.14 | 0.44 | 118 | 32_F_Novel | Coch | 4.53 | 12.19 | 7.64 | 8.13 | 2.09 | 0.26 |
| 54 | 17_Tbr1_S1 | Irx4 | 0.09 | 1.19 | 0.76 | 0.77 | 0.24 | 0.31 | 119 | 38_FmidiON | Foxp2 | 1.00 | 3.50 | 1.94 | 1.99 | 0.62 | 0.31 |
| 55 | 36_Novel | Coch | 2.20 | 8.27 | 3.25 | 3.74 | 1.50 | 0.40 | 120 | 38_FmidiON | Anxa3 | 0.25 | 1.60 | 0.74 | 0.78 | 0.36 | 0.46 |
| 56 | 36_Novel | Stxbp6 | 0.46 | 3.52 | 1.31 | 1.66 | 0.82 | 0.50 | 121 | 4_FminiOFF | Foxp2 | 1.74 | 3.20 | 2.42 | 2.35 | 0.37 | 0.16 |
| 57 | 12_ooDS_NT | Neurod2 | 0.61 | 1.71 | 1.13 | 1.15 | 0.23 | 0.20 | 122 | 4_FminiOFF | Gria1 | 1.53 | 3.06 | 2.19 | 2.16 | 0.42 | 0.19 |
| 58 | 12_ooDS_NT | Satb2 | 0.27 | 0.66 | 0.43 | 0.45 | 0.11 | 0.23 | 123 | 28_FmidiOFF | Foxp2 | 0.54 | 3.65 | 2.21 | 2.16 | 0.80 | 0.37 |
| 59 | 12_ooDS_NT | Mmp17 | 0.70 | 1.83 | 1.26 | 1.24 | 0.25 | 0.20 | 124 | 28_FmidiOFF | Sema5a | 0.57 | 2.34 | 1.44 | 1.48 | 0.43 | 0.29 |
| 60 | 35_Novel | Neurod2 | 0.00 | 2.39 | 1.42 | 1.46 | 0.58 | 0.40 | 125 | 3_FminiON | Foxp2 | 1.22 | 3.56 | 2.48 | 2.46 | 0.54 | 0.22 |
| 61 | 35_Novel | Satb2 | 0.00 | 2.94 | 0.84 | 0.90 | 0.55 | 0.62 | 126 | 3_FminiON | Irx4 | 0.98 | 1.97 | 1.59 | 1.58 | 0.22 | 0.14 |
| 62 | 35_Novel | Chrm2 | 0.08 | 1.06 | 0.56 | 0.53 | 0.27 | 0.52 | 127 | 44_Novel | Bhlhe22 | 0.00 | 0.53 | 0.21 | 0.22 | 0.17 | 0.80 |
| 63 | 29_Novel | Neurod2 | 0.00 | 1.56 | 0.99 | 0.87 | 0.34 | 0.39 | 128 | 44_Novel | Fxyd6 | 0.00 | 3.03 | 0.06 | 0.24 | 0.68 | 2.85 |
| 64 | 29_Novel | Satb2 | 0.42 | 3.53 | 0.88 | 1.04 | 0.70 | 0.67 | 129 | 11_Novel | Cacna1e | 0.00 | 1.29 | 0.72 | 0.74 | 0.30 | 0.41 |
| 65 | 29_Novel | Cpne5 | 0.71 | 3.08 | 2.16 | 2.00 | 0.76 | 0.38 | 130 | 11_Novel | Gm17750 | 0.46 | 2.23 | 1.24 | 1.28 | 0.37 | 0.29 |
